# Mechanisms Underlying the Complex Dynamics of Temperature Entrainment by a Circadian Clock

**DOI:** 10.1101/2021.04.28.441752

**Authors:** Philipp Burt, Saskia Grabe, Cornelia Madeti, Abhishek Upadhyay, Martha Merrow, Till Roenneberg, Hanspeter Herzel, Christoph Schmal

## Abstract

Autonomously oscillating circadian clocks resonate with daily environmental (zeitgeber) rhythms to organize physiology around the solar day. While entrainment properties and mechanisms have been studied widely and in great detail for light-dark cycles, entrainment to daily temperature rhythms remains poorly understood despite that they are potent zeitgebers.

Here we investigate the entrainment of the chronobiological model organism *Neurospora crassa*, subject to thermocycles of different periods and fractions of warm versus cold phases, mimicking seasonal variations. Depending on the properties of these thermocycles, regularly entrained rhythms, period-doubling (frequency demultiplication) but also irregular aperiodic behavior occurs. We demonstrate that the complex nonlinear phenomena of experimentally observed entrainment dynamics can be understood by molecular mathematical modeling.

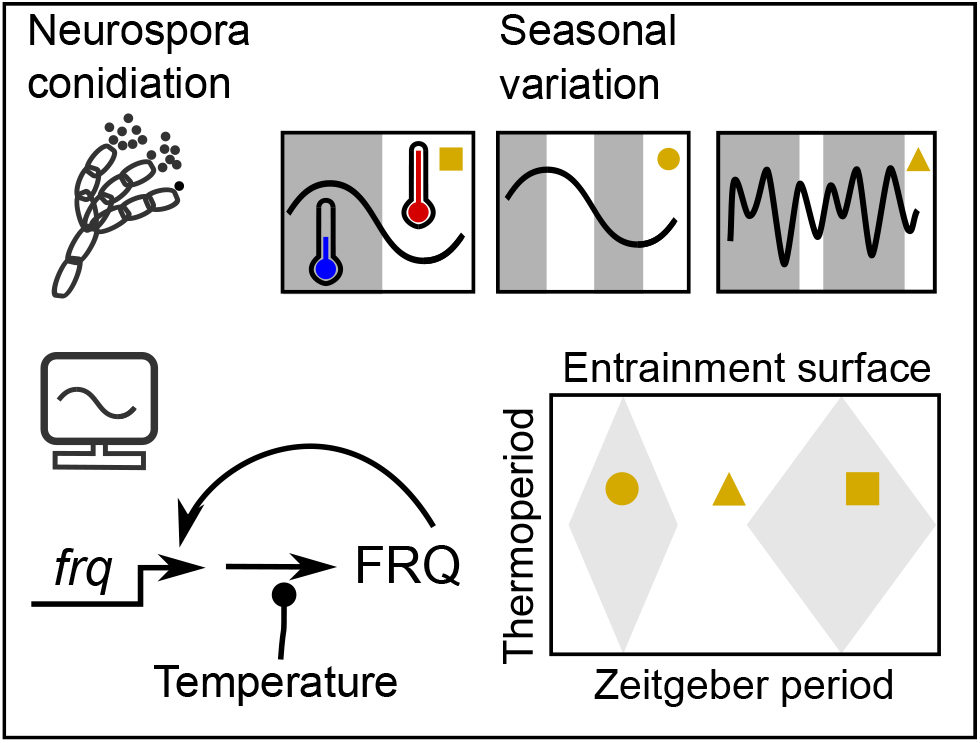

## 1 Introduction

From its outset, life on Earth is confronted with rhythmic changes in environmental conditions such as day and night, tides or seasons. In order to cope with daily changes of light and temperature, many organisms have evolved an autonomous circadian clock that rhythmically controls both behavior and physiology. A fundamental property of circadian clocks is that under physiological conditions, they entrain to an external zeitgeber such as light, food intake or temperature. By this means, physiological processes such as immune responses, metabolic activity or behavior are aligned rhythmically across the day. Thus, properties of zeitgeber cycles drive the evolution of the entire circadian system and it has been shown that geophysical differences in light conditions drive genetically encoded latitudinal clines of clock properties (Rosato et al. [1996], Sawyer et al. [1997], Hut et al. [2013]). In humans, misalignment between the clock and the external environment has been associated with cancer, diabetes and inflammatory diseases (Fuhr et al. [2015]). In plants, it leads to a decrease in photosynthesis and growth (Dodd et al. [2005]). The intricate regulation of the clock and its modulation by external stimuli requires a thorough understanding of generic clock principles. A rhythmic zeitgeber together with the intrinsic circadian clock it affects, constitutes a system of coupled periodic oscillators (Balanov et al. [2009]). Oscillator theory and mathematical modeling have proven to be useful tools for the investigation of circadian rhythms as they allow for a systematic study of entrainment parameters such as the occurrence of synchronization, entrainment amplitudes or entrainment phases *ψ* in dependence on the underlying zeitgeber and clock properties (Wever [1979]). For example, a prediction is that relatively short intrinsic periods *τ* lead to early entrainment phases (Wever [1965], Pittendrigh and Daan [1976]).

The filamentous fungus *Neurospora crassa* has been widely used as a circadian model organism in the past decades (Hurley et al. [2016]). *Neurospora crassa* has a fully sequenced genome and due to its haploid life cycle it is possible to perform genetic screens without backcrossing. In its asexual life cycle *Neurospora* produces visible spores, so-called conidia, which can be quantified by densitometric analysis (Belden et al. [2007]). Pioneered by Goldbeter and Ruoff the transcriptional feedback loop that leads to circadian rhythms of *Neurospora* has been modeled mathematically (Leloup et al. [1999], Ruoff et al. [2001], Francois [2005]). In particular, the role of protein degradation and mechanisms of temperature and glucose compensation have been studied in detail (Ruoff et al. [2005], Dovzhenok et al. [2015], Upadhyay et al. [2019]). More recently, oscillator theory has been applied to explore entrainment by light of different *Neurospora* strains (Rémi et al. [2010], Bordyugov et al. [2015]).

On the molecular level, self-sustained rhythms in *Neurospora* are generated by a delayed negative feedback loop with the frequency protein FRQ as the key element. The cycle starts with the transcription of the *frequency* gene (*frq*) driven by the heterodimer WCC, composed of the subunits WC-1 and WC-2. After a delay that is controlled by multiple phosphorylations and complex formations, FRQ protein inhibits its own transcription, thus closing the negative feedback loop which ultimately leads to autonomous circadian rhythmicity (Aronson et al. [1994], Dunlap [1999], Lee et al. [2000], Schafmeier et al. [2005], Upadhyay et al. [2020]). Notably, while light entrainment has been studied in *Neurospora* in detail both on the behavioral and molecular level, entrainment through temperature changes has received less attention (Rémi et al. [2010], Crosthwaite et al. [1995], Liu et al. [1997, 1998], Merrow et al. [1999], Rensing and Ruoff [2002], Lee et al. [2003]).

Here, we systematically explore the entrainment behavior of three different *Neurospora* strains (one wild type and two *frq* mutants), exposed to temperature cycles of up to five different lengths (*T* cycles). The combination of *T* cycles with a wild type strain, a short period mutant and a long period mutant created a dense network of *τ*-*T* relationships which would allow a granular understanding of the relationship of endogenous period to phase of entrainment. Furthermore, we varied the relative duration of the warm phase (27°C) with respect to the cold phase (22°C) of a zeitgeber cycle, ranging from 16% to 84% in 9 steps. In this way, we mimicked photoperiods with temperature cycles, further probing entrainment with various temporal structures. Alterations in zeitgeber structure can change the effective zeitgeber strength. Thus, the experimental design invites a thorough analysis of entrainment.

Using a combination of experiment and mathematical modeling, we address the following questions: What is the range of entrainment under varying conditions? Are there nonlinear phenomena outside of the 1:1 entrainment range? Do we find the expected association of the mismatch between *τ* and *T* and phase of entrainment (*ψ*)? How do seasonal environmental variations affect entrainment?

We find an astonishing complexity within the experimental recordings including 1:2 entrainment, superposition of two frequencies (tori), and reproducible aperiodic behavior that resembles deterministic chaos. Moreover, using a simple biophysically motivated molecular model of the temperature entrained *Neurospora crassa* core clock, we show that almost all observations including the aperiodic behavior can be explained by a single periodically driven oscillator.

## 2 Results

### 2.1 *Neurospora crassa* shows complex patterns of conidiation rhythms upon temperature entrainment

Circadian rhythms are generated at the single cell level through gene regulatory networks. Quantification of gene expression oscillations can be labor intensive and costly. Thus, more indirect measures from circadian clock controlled processes such as stem elongation and leaf movement rhythms in plants (Mayer [1966]) or behavioral activity rhythms in mammals (Johnson [1926]) and flies (Konopka and Benzer [1971]) are commonly used to deduce properties of the underlying circadian clock, such as the free-running period *τ* or phase of entrainment *ψ*. Here, we use densitometrically analyzed patterns of *N. crassa* conidia formation on agar medium, plated in glass tubes ('race tube assays'), as a proxy for circadian clock properties. In order to investigate mechanisms of temperature entrainment, we systematically exposed different strains of *N. crassa* to thermocycles of three different *T* cycles, namely *T* = 16h, 22h, and 26h. All experiments were constructed using cycles of 22°C and 27°C which corresponds to a zeitgeber amplitude of 2.5°C. In each series of *T* cycles, nine different thermoperiods namely *ϰ_T_* = 0.16, 0.25, 0.33, 0.4, 0.5, 0.6, 0.67, 0.75, 0.84 were tested. Thermoperiods are defined as the fraction 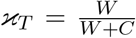 of the cycle length occupied by the warm phase *W*, where the cold phase is *C = T* − *W*. In *T* = 12h and *T* = 24h, entrainment was investigated at a reduced set of three different thermoperiods *ϰ_T_* = 0.25, 0.5, 0.75 and *ϰ_T_* = 0.16, 0.25, 0.33, respectively.

Qualitative changes in the dynamical behavior of conidiation rhythms occurred for variations in zeitgeber structures and endogenous period according to the underlying *N. crassa* strain (Fig. 1). For example, the wild type (*frq* ^+^) strain with a free-running period of *τ* ≈ 22h in constant conditions robustly entrains to an equinoctial thermocycle (*ϰ_T_* = 0.5) of period *T* = 22h that closely matches the period of the internal clock (Fig. 1 A). Transient dynamics approaching the entrained state (see first two days of Fig. 1 A) are indicative of entrainment effect as opposed to a simple coincidence between free-running and zeitgeber periods. Instantaneous phases, estimated by a Hilbert transformation, suggest entrainment, i.e. the conidiation cycle progresses by 2*π* (or 360°) along a full zeitgeber cycle, see (Fig. 1 B). Dominant peaks close to the zeitgeber period in the corresponding Lomb-Scargle spectrograms suggest frequency locking between the thermocycle and conidiation rhythms (Fig. 1 C).

**Figure 1:**
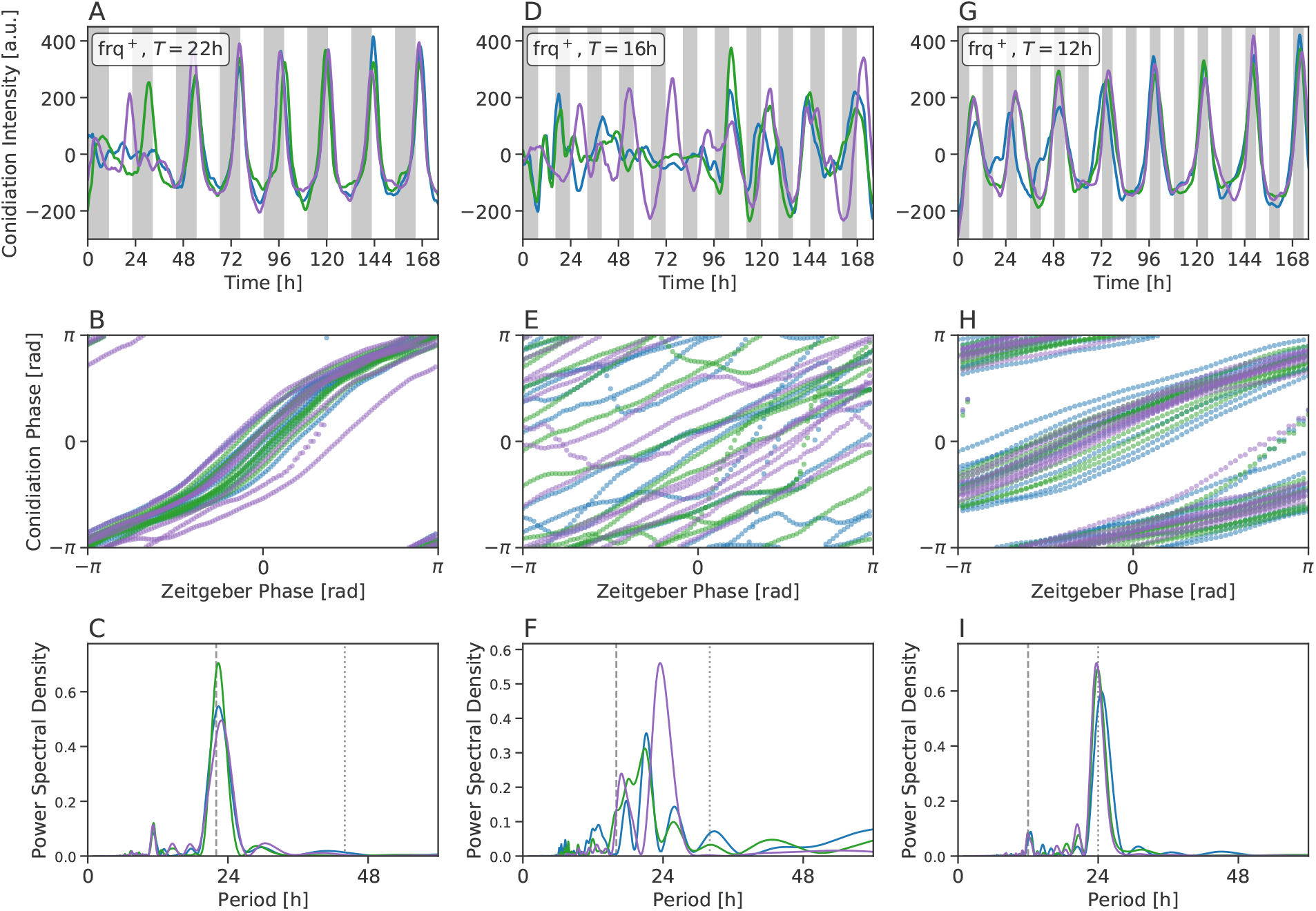
Complex dynamics upon temperature entrainment. A-C) Robust 1:1 entrainment can be observed for the *frq*^+^ strain under temperature cycles of *T* = 22h with 50% warm phase (*ϰ_T_* = 0.5). This is reflected in the densitometrically quantified time traces of conidiation patterns (A), in the phase locked dynamics between the zeitgeber and conidiation cycle (B) and in the spectrum of the corresponding oscillations as quantified by a Lomb-Scargle periodogram (C). D-F) Conidiation patterns of the *frq*^+^ strain do not entrain to a *T* = 16h thermocycle of *ϰ_T_* = 0.5. G-I) Frequency demultiplication or 2:1 entrainment can be observed in rhythmic conidiation patterns of the *frq*^+^ strain, subject to *T* = 12h thermocycles of thermoperiod *ϰ_T_* = 0.5. Different colors in panels (A)-(I) indicate biological replicates under the same experimental condition. Gray bars denote cold phases (22°C) while white background indicates warm phases (27°C).

Conidiation patterns fail to entrain if the period of an equinoctial (*ϰ_T_* = 0.5) thermocycle is reduced to *T* = 16h, 6h shorter than the *frq*^+^ strain’s internal free-running period of *τ* ≈ 22h (Fig. 1 D). No systematic dependency between the instantaneous phases of the zeitgeber cycles and the conidiation patterns was observed (Fig. 1 E). Lack of entrainment is further reflected in the corresponding Lomb-Scargle periodogram (Fig. 1 F), showing no agreement between the dominant spectral peaks and the entrainment period *T*.

A further reduction of the *T* cycle to *T* = 12h, close to half of the *frq*^+^ strains internal free running period, again leads to robust entrainment (Fig. 1 G and H). Here, one cycle of conidiation occurs every second full thermocycle, leading to a conidiation period of approximately 2*T* = 24h, see Fig. 1 I, a phenomenon, commonly known as frequency demultiplication (Bruce [1960], Merrow et al. [1999], Roenneberg et al. [2005]) or 1:2 entrainment (Pikovsky et al. [2001]).

We classified experimentally observed dynamics under temperature entrainment for all 99 different combinations of the three different strains, five different entrainment periods and up to nine different thermoperiods, using a semi-automated approach (Fig. 2). The observed dynamics are grouped into four categories, namely non-entrained, 1:1, 1:2 and 2:3 entrained oscillations (compare Fig. S1-S9).

**Figure 2:**
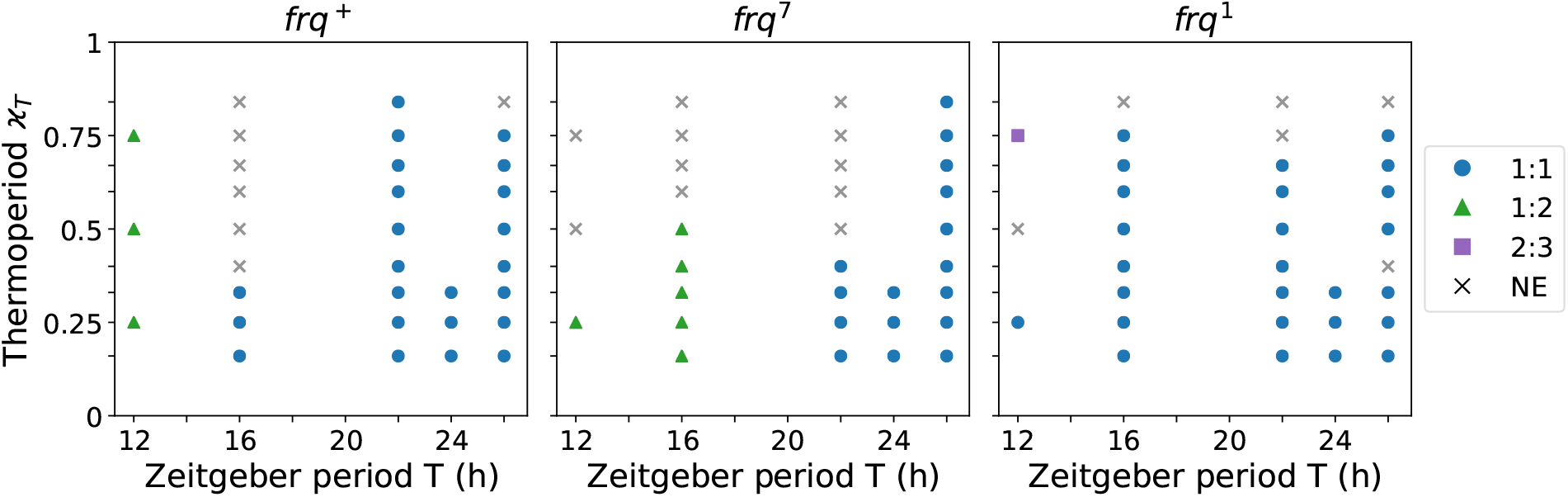
Circadian surface for temperature entrainment. Overview on *Neurospora crassa* conidiation experiments for wild type *frq*^+^ (left), long period mutant *frq*^7^ (middle), and short period mutant *frq*^1^ (right) strains, exposed to varying temperature cycle length (*T*; x-axis) and thermoperiods (*ϰ_T_*; y-axis). Each data point represents one experimental condition. Occurence of entrainment has been analyzed by inspecting raw data, phase plots and spectral decomposition via Lomb-Scargle periodograms, see Fig. S1 - S9. 1:1 entrainment is represented by cycles, 1:2 entrainment by triangles, 2:3 entrainment by squares, and conditions where no entrainment (NE) occurs by crosses.

### 2.2 Modeling temperature entrainment

In order to investigate the underlying principles of the experimentally observed complexity in temperature entrainment, we exploit a detailed mathematical model of the *Neurospora crassa* core clock as previously published (Hong et al. [2008]; *Hong Model*). The *Hong model* has been curated to explain rhythmic oscillations of core clock genes *frq* and *wc-1* in the wild type (WT) and the two mutant strains (*frq*^+^, *frq*^1^ and *frq*^7^, respectively) in constant darkness. Figure 3 A shows a schematic representation of the regulatory network. It consists of seven ordinary differential equations (ODEs) and 19 parameters (Eq. (1) and Table (1) in *Materials and Methods* show the detailed model structure and its corresponding kinetic parameters).

**Figure 3:**
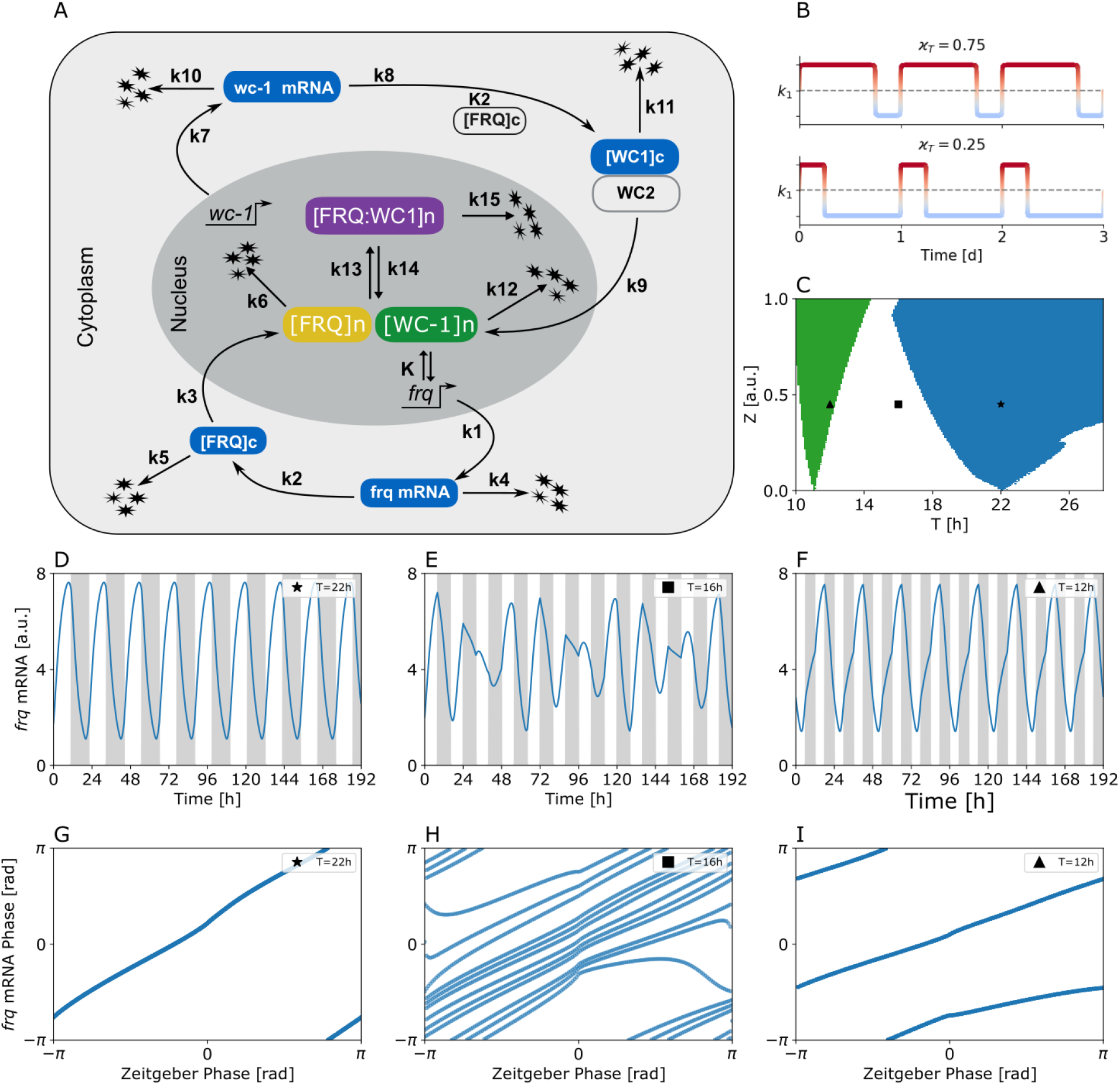
Different synchronization regimes (Arnold tongues) within a molecular clock model are able to explain the complex dynamics of Neurospora temperature entrainment. A) Schematic drawing of the regulatory network underlying the *Hong model*. B) Sketch of the temperature cycle induced regulation of the maximal transcription rate as described in *Materials and Methods* for two different thermoperiods *ϰ_T_*. For a given zeitgeber strength *z*_0_, the rate fluctuates between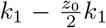 and 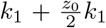 around its nominal value of *k*_1_. Phases of hot and cold temperatures are denoted by *red* and *blue* colors, respectively. C) 1:1 (*blue*) and 2:1 (*orange*; frequency demultiplication) synchronization regions (Arnold tongues) in the *Hong model*, entrained by equinoctial square wave temperature cycles as given by Eqs. (4, 5). D-F) Illustrative, representative example simulations of *frq* mRNA dynamics for equinoctial (*ϰ* = 0.5) temperature cycles at three different periods *T* = 22h, *T* = 16h, and *T* = 12h, respectively, at a constant zeitgeber strength of *z*_0_ = 0.45. G-I) Instantaneous phases of *frq* mRNA dynamics of sub-panels (D-F), plotted against corresponding zeitgeber phases. Parameter values corresponding to results plotted in sub-panels (D-I) have been highlighted by different markers in sub-panel (B), respectively.

While WC-1 and WC-2 have been shown to be involved in light entrainment (Froehlich et al. [2002], Liu [2003]), molecular details underlying temperature entrainment remain less understood. However, FRQ has been described to oscillate at a higher magnitude and amplitude with increasing temperatures (Liu et al. [1998]), potentially through alternative splicing of *frq* mRNA (Diernfellner et al. [2005]). Similar to other organisms, *Neurospora crassa* shows temperature compensation of its circadian free-running period (Gardner and Feldman [1981]). In order to reveal parameters that reproduce these features upon temperature impact, we applied a comprehensive sensitivity and bifurcation analysis of the *Hong model* (Figs. S10 and S11, respectively). As a result, changes in *frq* transcription and *frq* translation rates (parameters *k*_1_ and *k*_2_, respectively) are found to mimic temperature induced changes in amplitude while, the free-running period remains largely stable. Based on these qualitative observations, we constructed a conceptual model of temperature entrainment by assuming that temperature changes affect the clock by modulating the transcription rate *k*_1_ of *frq* mRNA.

### 2.3 Periodically driven limit cycle oscillator models explain complex entrainment behaviors

Using square-wave-like zeitgeber signals that approximate experimentally applied temperature cycles (see Equations (4) and (5) of *Materials and Methods* and Fig. 3 B), the *Hong model* shows typical hallmarks of entrainment (Pikovsky et al. [2001]). It can be easily entrained to zeitgeber signals with a period (*T*) close to the internal free-running period (*τ*). For increasing zeitgeber strength *z*_0_, the clock is able to entrain to a broader range of zeitgeber periods *T*, leading to tongue shaped entrainment region in the *z*_0_ − *T* parameter plane, called the *Arnold tongue* (Arnold [1987]), see *blue* area in Fig. 3 C. Within this tongue, 1:1 synchronization similar to the behavior of the *frq*^+^ strain under equinoctial temperature cycles of *T* = 22h occurs (Fig. 1 A,B and Fig. 3 D,G).

Entrainment not only occurs for zeitgeber periods close to the free-running period *τ*. Higher order synchronizaton is generally possible at rational fractions 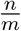 (n:m entrainment) where *n* is the number of cycles to which the internal clock adheres during *m* cycles of the zeitgeber signal (Balanov et al. [2009]), as illustrated for the 1:2 entrainment region (frequency demultiplication) of the *Hong model* in figure 3 C (*green*). Graphs F, and I correspond to dynamics observed for the *frq*^+^ strain under equinoctial temperature cycles of *T* = 12h (compare Fig. 1 G and H). The entrainment regions or *Arnold tongues* become smaller with increasing synchronization order (Fig. 3 C) as predicted from the literature.

No period or phase locking occurs outside the *Arnold* tongues. However, even in the absence of synchronization (absolute coordination), the zeitgeber is still able to affect the oscillations of the internal clock, a phenomenon called relative coordination (v. Holst [1938]). In this dynamical regime, complex phenomena leading to non-periodic dynamics (Fig. 3 E, H) as observed for the *frq*^+^ strain under equinoctial temperature cycles of *T* = 16h (compare Fig. 1 G and H) are characteristic.

Depending on zeitgeber strength (*z*_0_) and period (*T*) different qualitative behavior such as phase slipping, modulations, beating, quasi-periodicity or chaos can be underlying mechanisms of such non-periodic behavior (Fig. 4). Close to the borders of entrainment, right after leaving the entrainment regime, two types of slow modulations of the intrinsic oscillations typically emerge (Balanov et al. [2009], Granada et al. [2011]). At relatively low zeitgeber strength (*z*_0_) long-period amplitude modulations occur (Fig. 4 B *inset*). These are characterized by a torus in phase space (Fig. 4 B) whose stroboscopic map, i.e. the points extracted at the interval of the entrainment period *T*, forms a closed curve (Fig. 4 B *orange dots*) and a spectrum containing a peak at the period of the zeitgeber signal, a peak at the internal period of the circadian clock as well as peaks at combinations of these two periods (Fig. S12, Balanov et al. [2009], Granada et al. [2011]). As one gets closer to the entrainment region, the modulation period (i.e. of the amplitude envelope) lengthens tremendously while the diameter of the stroboscopic map remains relatively constant (Fig. S12). At higher zeitgeber strength (*z*_0_) beating is observed, i.e. the superposition of two frequencies, again characterized by a torus in phase space and a closed curve for the stroboscopic map (4 C). Characteristically, the beating period lengthens, the diameter of the stroboscopic map reduces and the spectral peak associated with the internal circadian period becomes smaller as one approaches the limits of entrainment (Fig. S13, Balanov et al. [2009]).

**Figure 4:**
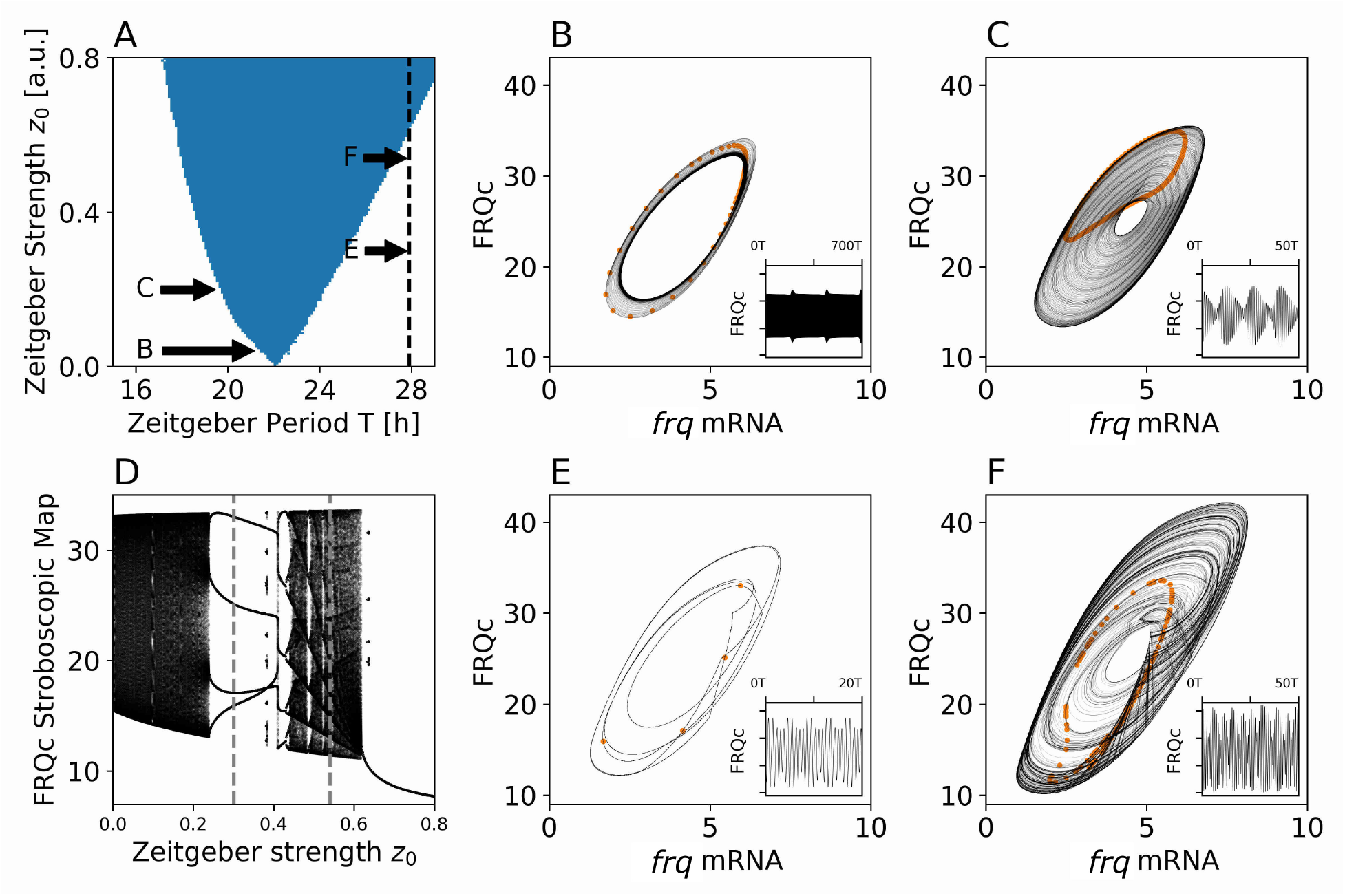
Complex dynamical behavior is observed outside the entrainment range. A) 1:1 synchronization region (*Arnold tongue*, *blue*) in the *Hong model*, entrained by a thermoperiod of *ϰ_T_* = 0.75. Combinations of *z*_0_ and *T* underlying panels B,C,E and F are indicated by arrows. Range of *z*_0_ at *T* = 27.9h underlying panel D is depicted by a dashed vertical line. B) Projection of a trajectory from the *Hong model* showing slow amplitude modulations for zeitgeber strength *z*_0_ = 0.04 and period *T* = 21.37h in the *frq* mRNA and cytosolic FRQ protein (FRQc) plane, after decay of transient dynamics. *Orange dots* show the associated stroboscopic map. *Inset* shows corresponding FRQc dynamics over the time of 700 entrainment periods *T*. C) Same as (B) showing beating behavior for *z*_0_ = 0.2 and *T* = 19.58h. D) One-parameter diagram of values from the FRQc stroboscopic map versus different zeitgeber strength *z*_0_. E) Same as (B) showing n:m higher order synchronization behavior for *z*_0_ = 0.3 and *T* = 27.9h. F) Same as (B) showing chaotic behavior behavior for *z*_0_ = 0.54 and *T* = 27.9h.

By systematically investigating the dynamics for different zeitgeber strength (*z*_0_) at a given period (*T*) as characterized by the stroboscopic map of cytosolic FRQ protein (*FRQc*) oscillations, a collection of qualitatively different dynamics can be observed (Fig. 4 D-F). With increasing zeitgeber strength (*z*_0_) at *T* = 27.9h, the system evolves from quasiperiodic torus oscillations through a couple of higher order synchronization regimes (e.g. Fig. 4 E) and chaotic behaviors (e.g. Fig. 4 F) until it reaches the 1:1 entrainment region (*Arnold tongue*) at *z*_0_ ≈ 0.62 (compare also Fig. S14 for an illustration).

In summary, fundamental synchronization properties of a limit cycle oscillator such as the *Hong model*, when entrained to a periodic forcing signal, explain the complex patterns of conidiation rhythms that have been observed under equinoctial (*ϰ* = 0.5) temperature entrainment in the wild type *Neurospora crassa* strain (compare similarities in Fig. 1 and 3). In the next section we extend this analysis to the full circadian surface, i.e. all combinations of zeitgeber periods *T* and thermoperiods *ϰ_T_* as summarized in figure 2 A.

### 2.4 Simulated clock dynamics are able to explain behavior across the full circadian surface of the *frq*^+^ wild type strain

Temperature entrained conidiation rhythms of the wild type *Neurospora crassa* strain (*frq*^+^) show different dynamical regimes, depending on zeitgeber period (*T*) and thermoperiod (*ϰ_T_*; Fig. 2 A). 1:1 entrainment is observed when *T* is close to *τ* ≈ 22h, 1:2 entrainment occurs when *T* is around 50% of *τ*, and non-entrained dynamics occur when *T* = 16h and *T* = 26h. Especially for short zeitgeber periods of *T* = 16h, the *frq*^+^ strain entrains better under short thermoperiods at the experimentally chosen zeitgeber amplitude.

For an arbitrarily chosen zeitgeber strength *z*_0_, most of the simulated clock dynamics (~ 50% in Fig. 5 A) do not agree with experimentally observed conidiation rhythms. The location, width, and tilt of the *Arnold onions* (synchronization regimes, Schmal et al. [2015]) is determined by both properties of the internal clock and the external zeitgeber. While the width of the Arnold onion is determined by multiple factors such as, e.g., the strength of the zeitgeber signal or the amplitude and radial relaxation rate of the circadian clock, the position of the entrainment regions in the *ϰ_T_* - *T* plane is mainly determined by the clock’s internal free-running period *τ*. In autonomous dynamical systems such as the *Hong model*, the internal free-running period *τ* can always be rescaled, see *Materials and Methods*. Along these lines, we searched for minimal modifications of our conceptual temperature entrainment model to better mimic experimental findings. To this end, we exhaustively investigated different combinations of internal periods *τ* and external zeitgeber strengths *z*_0_ for their fit of the data (Fig. 5 B). We fitted the entrainment state of the data for each of the 33 combinations of zeitgeber period *T* and thermoperiod *ϰ_T_*, i.e. we asked whether a non-entrained, 1:1 entrained or 1:2 entrained dynamics was observed and compared this with the dynamics described in figure 2.

**Figure 5:**
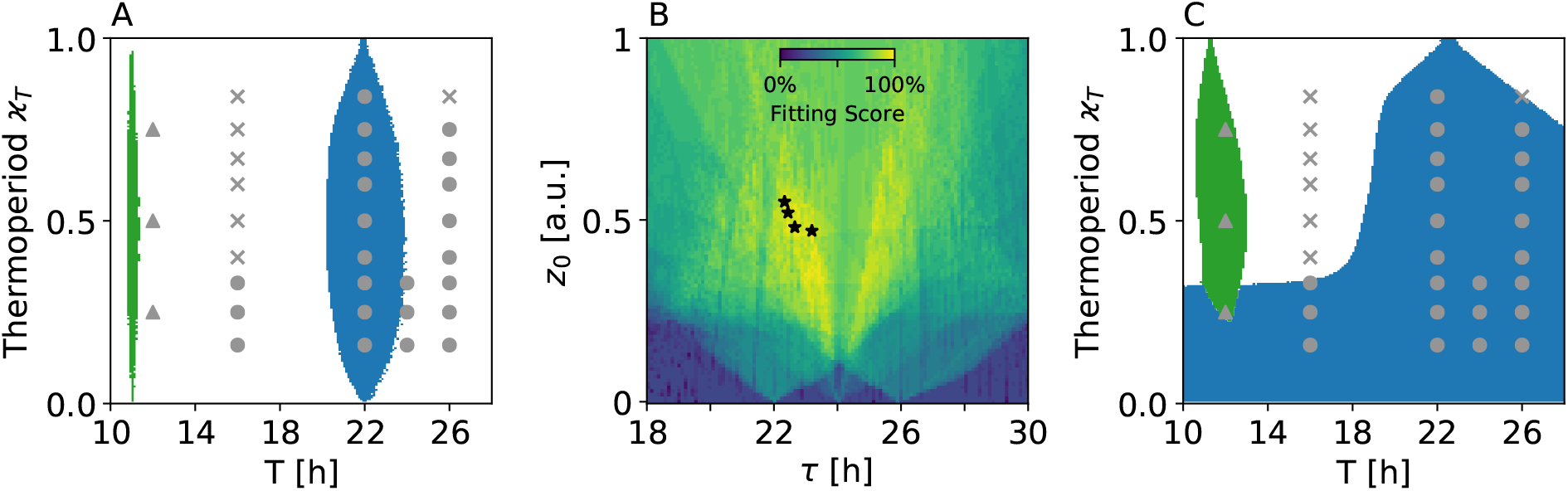
Good agreement of experimental and simulated wild type (*frq*^+^) Neurospora crassa entrainment dynamics after proper scaling of the models free running period and zeitgeber strength. A) Experimentally observed dynamics (gray symbols) together with simulated 1:1 (*blue*) and 2:1 (*green*; frequency demultiplication) synchronization regions in the thermoperiod - zeitgeber period parameter plane (Arnold onions). A zeitgeber strength of *z*_0_ = 0.1 and the nominal parameter set of the *Hong model* (see Table 1) have been used. B) Fitting scores for different zeitgeber strength *z*_0_ and scaled internal free-running periods *τ*. The fitting score in percent denotes the fraction of simulated dynamics that qualitatively matches the experimentally observed ones, i.e. whether 1:1 entrainment, 2:1 frequency-demultiplication or unentrained dynamics are observed. *Stars* denote parameter combinations that lead to fitting score of 100%. Experimentally observed entrainment dynamics and simulated synchronization regions (Arnold onions) as in panel (A) for an exemplary “optimal” parameter combination of panel (B), namely *z*_0_ = 0.47 and *τ* ≈ 23.19h.

**Table 1:** Kinetic parameters of the *Hong model* as published in (Hong et al. [2008]).

| Description | Parameter | Value ($\text{h}^{-1}$) |
| --- | --- | --- |
| *frq* overexpression | $k_{01}$ | 0.00 |
| basal *wc-1* mRNA translation | $k_{02}$ | 0.00 |
| *frq* transcription | $k_1$ | 1.80 |
| *frq* mRNA translation | $k_2$ | 1.80 |
| import to nucleus | $k_3$ | 0.05 |
| *frq* mRNA degradation | $k_4$ | 0.23 |
| degradation | $k_5$ | 0.27 |
| degradation | $k_6$ | 0.07 |
| *wc-1* transcription | $k_7$ | 0.16 |
| *wc-1* mRNA translation ( dependent) | $k_8$ | 0.80 |
| import to nucleus | $k_9$ | 40.00 |
| *wc-1* mRNA degradation | $k_{10}$ | 0.10 |
| degradation | $k_{11}$ | 0.05 |
| degradation | $k_{12}$ | 0.02 |
| complex formation | $k_{13}$ | 50.00 |
| complex dissociation | $k_{14}$ | 1.00 |
| complex degradation | $k_{15}$ | 5.00 |
| binding to *frq*-promoter | $K$ | 1.25 |
| mRNA complex formation | $K_2$ | 1.00 |

It turns out that scaling of free-running *τ* and zeitgeber strength *z*_0_ is sufficient for an “optimal” fit, i.e. all entrainment states of the experiment can be reproduced by simulation (Fig. 5 B and C). In order to mimic the experimentally observed 1:2 entrainment at *T* = 12h, slightly longer values of *τ*_opt_ in comparison to the 22h free-running period under constant darkness of the *Hong model* emerge (Fig. 5 B; *stars*). Interestingly, our fit leads to values that are in accordance with the experimentally observed free-running period and inter-individual period heterogeneity (Diegmann et al. [2010]).

### 2.5 Damped oscillations at cold temperatures facilitate entrainment under short thermoperiods

At zeitgeber period *T* = 16h, experimental conidiation rhythms seemed to entrain at short thermoperiods for *ϰ_T_* = 0.16, 0.25, 0.33 but failed to entrain at longer thermoperiods. This behavior is not reproducible with the onion shaped entrainment region of a weakly forced circadian system as shown in figure 5 A. Instead, “optimal” fits lead to a 1:1 entrainment region that widens under short thermoperiods. This widening can be explained by the bifurcation structure of the *Hong model* (Fig. S11, parameter *k*_1_). As zeitgeber amplitudes get large, the *Hong model* eventually enters two qualitatively different dynamical regimes under cold and warm temperatures. While it still behaves as a self-sustained oscillator under warm temperatures, it shows slightly damped oscillations under cold conditions (reverse *Hopf* bifurcation). Damped oscillators typically synchronize more easily by an external force in comparison to self-sustained pacemakers. This facilitated entrainment under cold temperatures leads to a broadening of the Arnold onion towards shorter thermoperiods in the *Hong model*, as illustrated for *z*_0_ = 0.47 in figure 5 C. A similar behavior has been recently observed for a light-entrained *Goodwin* oscillator under long photoperiods (Ananthasubramaniam et al. [2020]).

### 2.6 Predicted correlations between *ψ*, *ϰ_T_* and *T* are supported by experiment

In the case of synchronization, a stable phase relation between the zeitgeber signal and the internal clock, commonly termed the phase of entrainment *ψ*, emerges. A proper phase of entrainment is of fundamental importance as it ensures that diurnal physiological processes are aligned at appropriate times around the day. Fig. 6 A shows simulated entrainment phases *ψ* of cytosolic FRQ protein (FRQc) oscillations at different thermoperiods *ϰ_T_* and two different zeitgeber periods *T* = 22h and *T* = 26h, corresponding to the phases along vertical cross-sections through figure 5 C. The two periods correspond to those experimentally investigated periods showing the largest fraction of 1:1 entrained conidiation rhythms across varying thermoperiods in the *frq*^+^ strain. Simulations predict later phases of entrainment *ψ* with increasing thermoperiods *ϰ_T_* and decreasing zeitgeber periods *T* (Fig. 6 A). For *T* = 26h the shift of *ψ* towards later phases with increasing *ϰ_T_* saturates and reaches a plateau after *ϰ_T_* ≳ 0.5. The dependencies of the entrainment phase *ψ* on *ϰ_T_* and *T* are qualitatively similar to the corresponding behavior in temperature entrained conidiation patterns. Analogous to simulated 1:1 entrained FRQc oscillations, conidiation rhythms of the *frq*^+^ strain show consistently later phases with decreasing zeitgeber period *T* and in most cases a monotonic increase of *ψ* with increasing thermoperiod *ϰ_T_* (Fig. 6 B-D).

**Figure 6:**
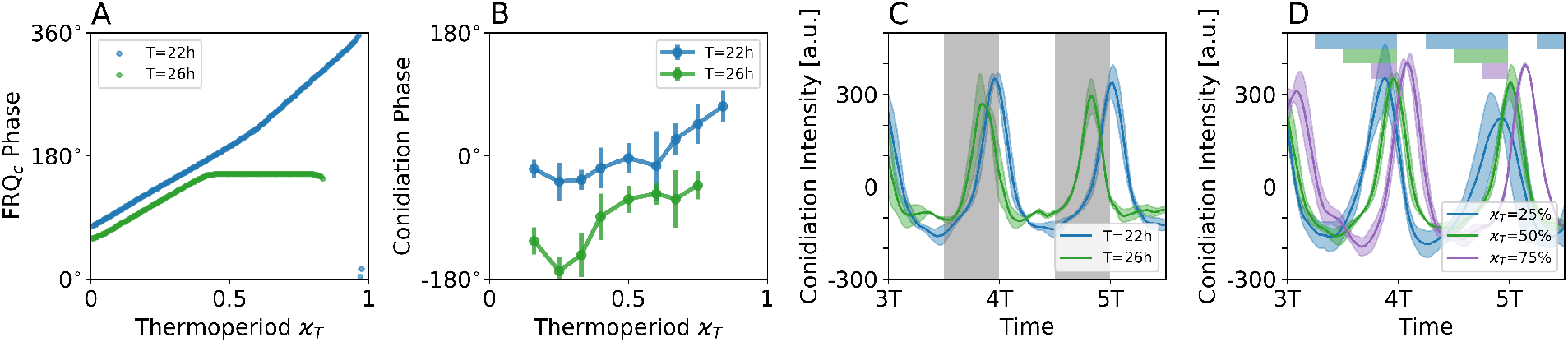
Earlier phases in longer entrainment cycles and later phases for increasing thermoperiods are observed in experiment and simulation. A) Simulated entrainment phases of the *Hong model* using parameters as in Fig. 5 C under different thermoperiods *ϰ_T_* and two different zeitgeber periods, namely *T* = 22h (*blue*) and *T* = 26h (*green*). B) Entrainment phases determined from 1:1 synchronized conidiation patters of the *frq*^+^ strain upon *T* = 22h (*blue*) and and *T* = 26h (*green*) entrainment cycles, using a peak-picking method. Each data point represents the circular mean and standard deviation, determined from all detected oscillation peaks (acrophases) of all three replicates at a given *ϰ_T_* and *T*. Peaks from the first three entrainment cycles have been neglected as transients. C) Exemplary 1:1 synchronized conidiation patterns under equinoctial temperature entrainment of period *T* = 22h (*blue*) and *T* = 26h (*green*) show the behavior as expected from simulations, namely earlier phases for longer entrainment cycles. Analogously to panel (C), exemplary 1:1 synchronized conidiation patterns under *T* = 22h show earlier phases for shorter thermoperiods *ϰ_T_* as expected from simulations of the *Hong model*. Gray shaded areas in panel C denote periods of cold temperatures. Colored bars at the top of panel D denote periods of cold temperatures under different thermoperiods. Bold lines on panels C-D denote averages while transparent areas denote standard deviations of all three conidiation pattern replicates of the *frq*^+^ strain as found in Fig. S1.

## 3 Discussion

Circadian entrainment has been predominantly studied using light as a synchronizing cue. Entrainment to temperature cycles has attracted less attention even though circadian clocks have been shown to synchronize to thermocycles of surprisingly low amplitudes such as 1 − 2°C in poikilotherms (Hoffmann [1969], Wheeler et al. [1993], Somers et al. [1998], Rensing and Ruoff [2002]) or tissue cultures of homeotherms (Brown et al. [2002], Abraham et al. [2010], Buhr et al. [2010]). In the present paper we find diverse and complex dynamical responses during the entrainment of circadian *Neurospora crassa* conidiation rhythms subject to temperature cycles and subsequently show that this complexity can be well understood by mathematical models of molecular regulatory networks underlying the intracellular circadian clock.

Entrainment of oscillatory systems generally depends on properties of the intrinsic oscillator as well as the periodic stimulation. We analyzed these differential contributions by systematically varying the period *T* and thermoperiod *ϰ_T_* in temperature entrainment experiments using three different *Neurospora crassa* strains with different intrinsic circadian periods *τ*. While the wild type *frq*^+^ and the long period *frq*^7^ strains consistently show regular 1:1 entrainment only for zeitgeber periods that are relatively close to their previously published intrinsic free running periods, short period mutant *frq*^1^ is generally able to entrain to a broader range of *T* -*ϰ_T_* combinations. Higher order 1:2 synchronization also known as frequency demultiplication can be observed at zeitgeber periods close to 50% of the strains intrinsic period ((Merrow et al. [1999]), Fig. 2). Such a behavior has been analogously shown for light-entrained conidiation rhythms of *Neurospora crassa* (Rémi et al. [2010]). Noteworthy, the wild type *frq*^+^ and long period *frq*^7^ strains generally entrain better to short thermoperiods in comparison to long thermoperiods, being in contrast to the previously theoretically predicted symmetrical behavior for conceptual amplitude-phase oscillator models (Schmal et al. [2015], Diekman and Bose [2018]). Outside the synchronization regimes, we observe differently pronounced non-entrained and irregular behavior, characterized by drifting phases with respect to zeitgeber signals or fluctuating instantaneous periods and amplitudes.

To study principles underlying the experimentally observed complex entrainment behaviors, we employed a previously published detailed ordinary-differential-equation based mathematical model (Hong et al. [2008]) that considers the main regulatory interactions of the *Neurospora crassa* core clock. By using a detailed sensitivity and bifurcation analysis of the model parameters, constrained by experimentally observed behaviors such as temperature compensation (Gardner and Feldman [1981]) and the higher baseline expression and oscillation amplitude of FRQ protein at higher temperatures (Liu et al. [1998]), we suggest variations in *frq* transcription and translation rates as a potential driver underlying temperature entrainment. While this approach differs from the common assumption that changes in temperature affect all kinetic parameters, often modeled via Arrhenius equations (Ruoff et al. [2005], Tseng et al. [2012]), it reduces the number of necessary parameters tremendously and thus facilitates the model analysis and interpretation.

Using variations in *frq* transcriptional rate as a mechanism of temperature entrainment, we varied the intrinsic free-running period (*τ*) and zeitgeber strength (*z*_0_) and qualitatively compared the simulated entrainment states with the experimentally observed dynamics. This is tantamount to fitting the synchronization regimes (i.e. *Arnold tongues* and *Arnold onions*) to experimental data in the zeitgeber period *T* and thermoperiod *ϰ_T_* parameter plane. Similar approaches have been previously used to understand, for example, entrainment behavior of cardiac cells (Guevara et al. [1981], Glass et al. [1984]) or photoperiodic entrainment of plant circadian clocks (De Caluwe et al. [2017]). In summary, the qualitative synchronization behavior can be reproduced for all experimentally tested combinations of *T* and *ϰ_T_* in the *frq*^+^ wild type strain. Our unbiased optimization approach led to free-running periods *τ*, close to those that have been experimentally observed. Thus, the dependencies between synchronization regimes and free-running periods as predicted from oscillator theory (Balanov et al. [2009], Pikovsky et al. [2001]) are reproduced by our experiments.

For conceptual (amplitude-phase) oscillator models, we have previously shown that entrainment regions in the zeitgeber period and photoperiod paramater plane adopt an onion-shaped geometry, called *Arnold onions* (Schmal et al. [2015, 2020]). We find the same geometrical structure for the entrainment region in the case of the detailed molecular *Hong model* (Fig. 5 A). The 1:1 synchronization region has its widest entrainment range at the equinox and tapers towards zeitgeber periods (*T*) that correspond to the circadian clock’s internal free-running period *τ* for increasingly extreme thermoperiods (*ϰ_T_* ≈ 0 or *ϰ_T_* ≈ 1). For photoperiodic entrainment, a tilt of this onion-shape geometry is expected, since free-running periods often differ under constant darkness and constant light, known as Aschoff’s rule (Aschoff [1958]). Typically, the Arnold onion tapers towards these different values of *T* under extremely long or short day-length (Schmal et al. [2015]). In the case of temperature entrainment, such a tilt is not expected due to the temperature compensation of free-running period *τ*, a behavior that is faithfully mimicked by the temperature entrained *Hong model* (Fig. 5 A, *blue*).

Our optimization approach predicts that the experimentally observed broader entrainment regime for short thermoperiods relies on the bifurcation structure of the *Neurospora crassa* circadian clock. Bifurcations are qualitative changes of systems dynamics due to changes in a certain parameter. Here, our optimization based modeling approach predicts that the ciradian clock of *Neurospora crassa* moves from a self-sustained to a damped oscillatory regime for increasingly cold temperatures which ultimately leads to a broad entrainment range under short thermoperiods as damped oscillators can be more easily entrained. Interestingly, this is in line with experimental observations, showing an increasing damping and lowering amplitudes among conidiation patterns for decreasing temperatures (Liu et al. [1997]).

Simulations of the modified *Hong model* subject to temperature cycles reveal qualitatively different dynamics between the synchronization regimes. Close to the limits of entrainment typically beating and modulations (tori) were observed, similar to results from conceptual amplitude-phase models (Balanov et al. [2009], Granada et al. [2011]). Between successive higher-order synchronization regimes, we identified regions of chaotic dynamics, analogous to what has been found in other molecular circadian clock models subject to light entrainment (Gonze and Goldbeter [2000], De Caluwe et al. [2017]). Such non-autonomous chaotic behavior resembles the experimentally observed irregular aperiodic conidiation patterns in the absence of synchronization (e.g. Fig. 1 D). However, it should be noted that the existence of chaos cannot be formally proven within the typical short time scales of circadian experiments (Bradley and Kantz [2015]).

While experimental observations can be reliably reproduced in case of the wild type *frq*^+^ strain, the dynamical behavior of the long period *frq*^7^ and short period *frq*^1^ mutant can only be reproduced in up to ≈ 82% of the T and *ϰ_T_* combinations (compare Fig. S15). However, simulated dynamics of the *frq*^7^ and *frq*^1^ strain have been obtained by parameter changes as described in (Hong et al. [2008]). Since such study focused on dynamics in constant darkness without considering entrainment characteristics, it would be interesting to further investigate in future studies whether other combinations of parameter changes could reproduce the constant darkness free-running periods and temperature entrainment dynamics at the same time.

In conclusion, our study reveals complex temperature entrainment behavior for *Neurospora crassa* conidiation rhythms including higher order synchronization and nonlinear phenomena such as chaos or quasiperiodicity. We reveal design principles that underlie this complexity by mathematical modeling. Our proposed methodology of fitting model parameters to synchronization features at different entrainment protocols (e.g. varying thermo- or photoperiods) can be easily adapted to other systems of externally driven biological oscillators.

## 4 Materials and methods

### 4.1 Genetic material

Three *Neurospora crassa* strains with different free running periods *τ*_DD_ in constant darkness were used in the present study: The standard laboratory strain *bd A 30-7* (*τ* = 22*h*, denoted as *frq* ^+^) the *frq* ^1^ strain (*τ* = 16*h*) and the *frq* ^7^ strain (*τ* = 29*h* and deficient in temperature compensation at low temperatures) (Sargent and Kaltenborn [1972], Loros et al. [1989]). Both *frq* ^1^ and *frq* ^7^ derive from the *frq*^+^ strain but additionally carry a mutation in the *frq* gene (Loros et al. [1989]). They can be obtained from the Fungal Genetics Stock Center (Kansas City, KS, USA).

### 4.2 Temperature cycles and race tube assays

Temperature cycles were generated in customized incubators containing two integrated waterbaths that facilitated precise alternations between warming and cooling. Temperatures were thereby varied between 22 °C and 27 °C and *Neurospora* conidiation was assessed on race tubes filled with 10 ml of molten race tube media (2% Agar, 1X Vogel’s solution, 0.5% Arginine, 1 ng/1 ml Biotin, no glucose). All tubes were inoculated and germinated for 24 h in constant light at 25 °C before exposure to temperature cycles in darkness. Growth rates were marked at intervals under red light. Conidiation was then measured using densitometric analysis as previously described (Rémi et al. [2010]).

### 4.3 Data analysis

*Neurospora* conidiation patterns were transformed and normalized to densitometric intensities. Instantaneous phases were computed using the `hilbert` function from the `scipy.signal` library in `python`. Spectral analysis was performed after removal of the first 48h which were considered transient dynamics using the `LombScargle` method from the `Astropy` package. In cases where conidiation stopped before the end of the experiment run time, possibly due to medium limitations, corresponding time points were excluded from downstream analysis. Data were categorized as entrained or non-entrained according to three different criteria commonly used in the analysis of nonlinear dynamcis, namely the critical inspection of 1) raw time-series 2) phase-portraits and 3) spectra (Bergé et al. [1984], Schuster [1994], Erzberger et al. [2013]).

### 4.4 Ordinary differential equations and kinetic parameters

The dynamical evolution of the *Neurospora crassa* core clock follows equations

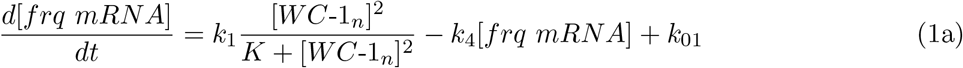

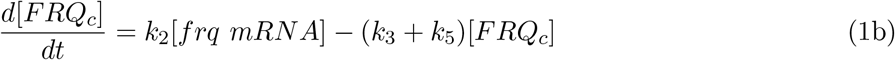

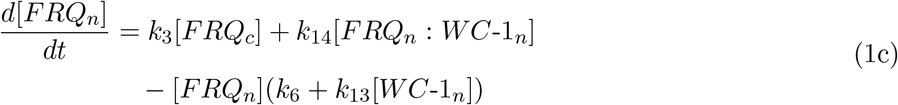

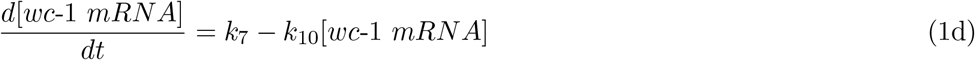

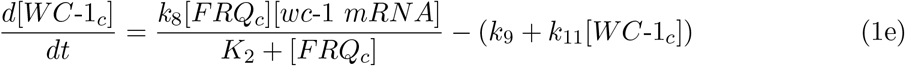

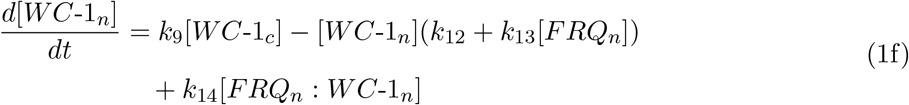

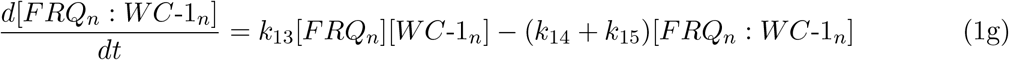

as previously published (Hong et al. [2008]), using the nominal parameter set as given in Table 1 for a subsequent optimization. Dynamics of the long period *frq*^7^ and the short period *frq*^1^ strain are simulated by setting *k*_5_ = 0.15h^−1^ and *k*_6_ = 0.01h^−1^ or *k*_3_ = 0.15h^−1^, *k*_5_ = 0.4h^−1^ and *k*_6_ = 0.1h^−1^, respectively, as described in (Hong et al. [2008]).

### 4.5 Re-scaling of intrinsic period

Using vector notation we can rewrite equations (1a)-(1g) as 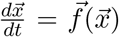 with 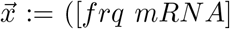 [*FRQ_c_*], [*FRQ_n_*], [*wc*-1 *mRNA*], [*WC*-1*_c_*], [*WC*-1*_n_*], [*FRQ_n_*: *WC*-1*_n_*])*^T^*. Such autonomous dynamical systems with 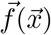 not explicitly depending on *t*, can always be rescaled in time. Defining a new variable *t′ = ct* with *c* being constant and using the chain rule

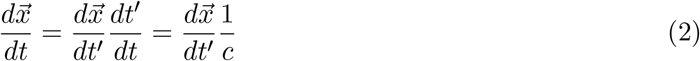

we can rewrite equations (1a)-(1g) as

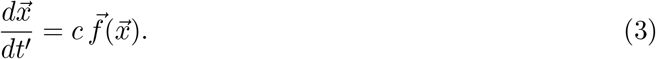

Solutions of this equation are identical to those obtained from (1a)-(1g) with the time axis being rescaled via a factor *c*. By this means we can freely choose the intrinsic free-running period *τ* under constant conditions as used during our optimization protocol underlying Fig. 5 B and Fig. S14 A,C.

### 4.6 Implementation of zeitgeber signal

To simulate entrainment in the Hong model, we chose a zeitgeber function with flexible period, thermoperiod and amplitude (zeitgeber strength), i.e.

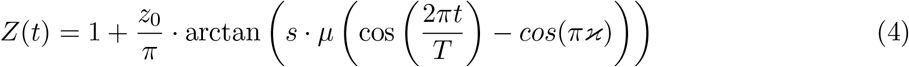

similar to what has been described in (Schmal et al. [2015]). Here, *z*_0_ is the zeitgeber strength, *s* is the steepness of the function, *T* is the period, *ϰ* is the thermoperiod and 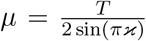 defines the slope at the switch points so that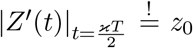 In this study we set *s* = 100. Based on our bifurcation analysis and on earlier works demonstrating the importance of *frq* transcription and translation for temperature entrainment (Liu et al. [1997, 1998]), we chose to modulate the *frq* transcription rate *k*_1_ to investigate entrainment in the *Hong model*. Note that *frq* transcription rate *k*_1_ and *frq* translation rate *k*_2_ show similar bifurcation patterns with a positive effect on the amplitude and little to no effect on the period and could thus both be used to implement entrainment from a theoretical perspective. Modulating the *frq* transcription rate *k*_1_ with *Z*(*t*) gives:

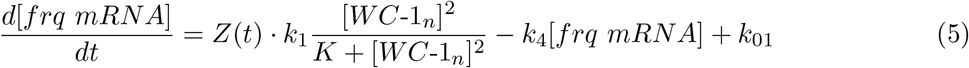

### 4.7 Numerical Simulations

Simulations were performed using the `scientific python` environment. Ordinary differential equations were solved using the odeint function from the `scipy.integrate` module with a time step of Δ*t* = 0.1. Simulations were performed for 200 zeitgeber cycles of which the first 100 cycles were excluded to avoid transient effects that might arise at the vicinity of the Arnold tongue.

Determination of the entrained state and order of synchronization within simulated dynamics has been determined as previously described (Schmal et al. [2015]). Bifurcation analysis was performed using XPP-Auto (Ermentrout [2002], Schmal et al. [2014]).

## Supplementary Material

**Figure S1:**
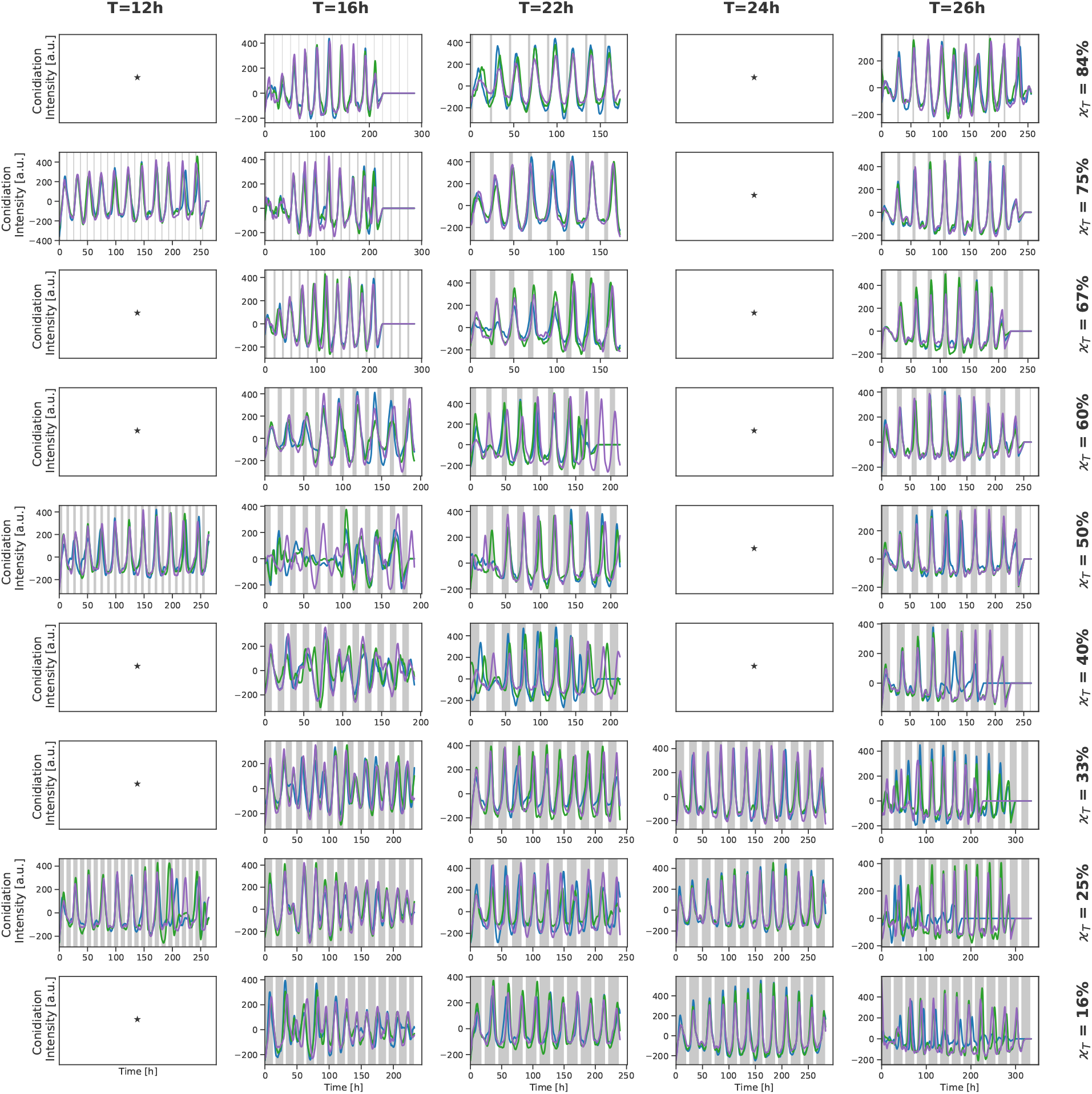
Experimental time series of the *frq^+^* (WT) strain. Conidiation patterns obtained by densitometric quantification. For each combination of Zeitgeber period (*T*) and thermoperiod (*ϰ_T_*), up to three race tube assays have been investigated. Differing colors denote for the individual replicates. Stars denote Zeitgeber period and thermoperiod combinations for which no data has been collected.

**Figure S2:**
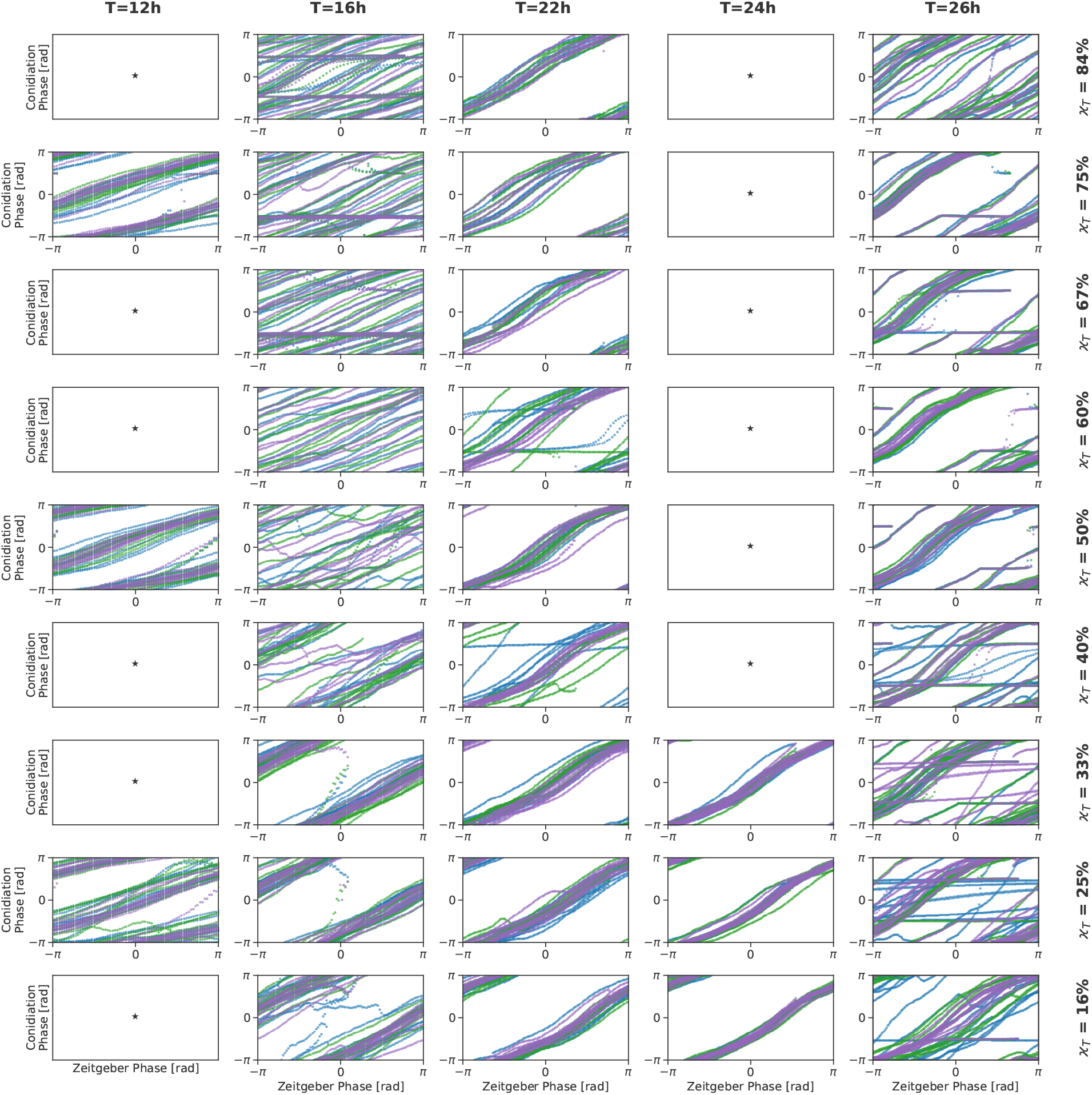
Phase plots for the *frq^+^* (WT) strain data. Instantaneous phases of the Zeitgeber cycle (x-axis) plotted against the instantaneous phases of the conidiation patterns (y-axis), determined from the experimental data as shown in Figure S1. After calculating instantaneous phases, the first 48h have been disregarded for plotting as potential transients during the entrainment process. Colors correspond to the the color coding in Figure S1.

**Figure S3:**
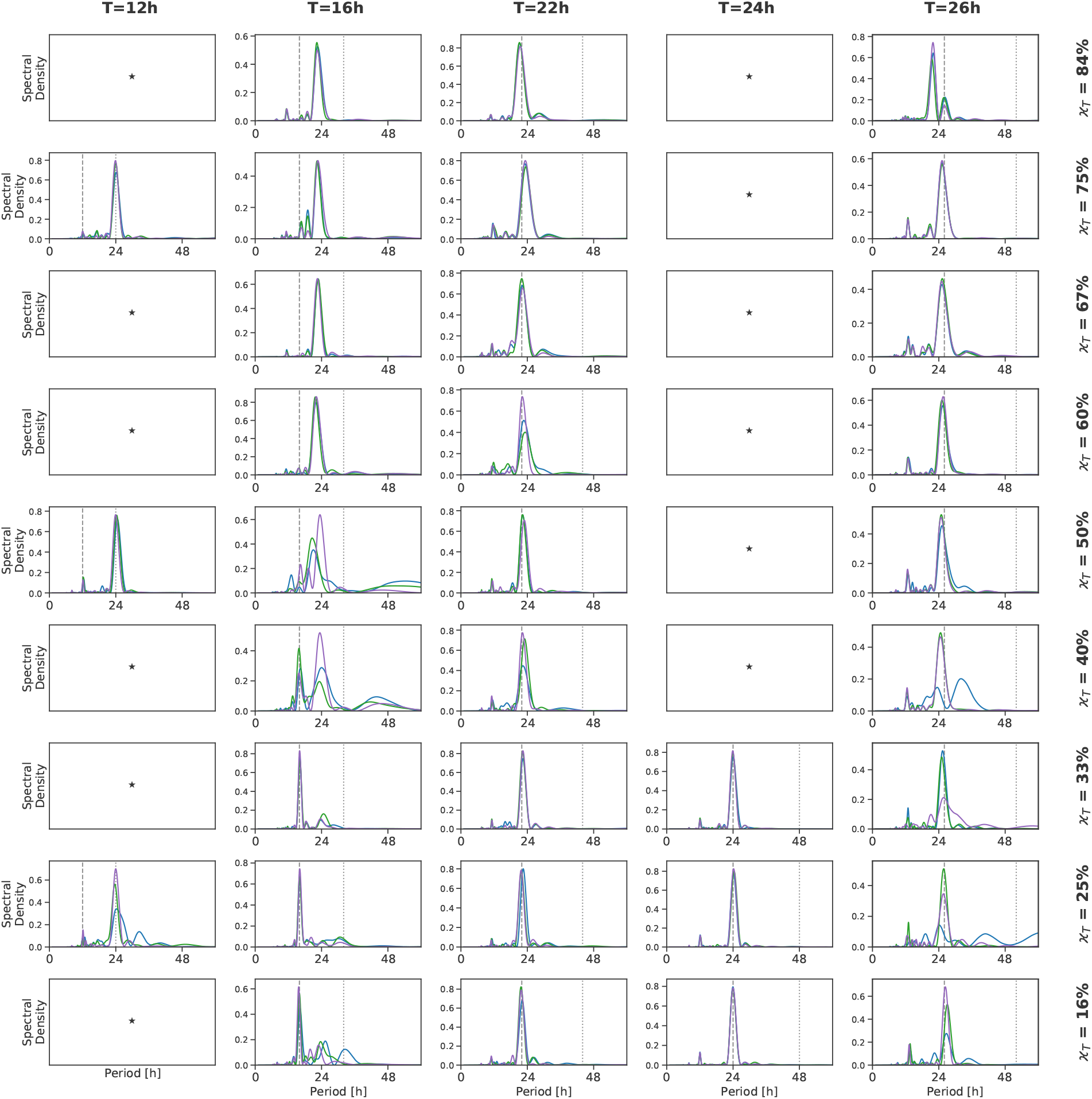
Lomb-Scargle periodograms for the *frq^+^* (WT) strain data. Lomb-Scargle periodograms, calculated for experimental data shown in Figure S1. The first 48h have been omitted from the analysis as transient dynamics during the entrainment process. Colors correspond to the color-coding in Figure S1. Dashed gray lines denote the Zeitgeber period *T* and dotted gray lines denote a period of 2*T*.

**Figure S4:**
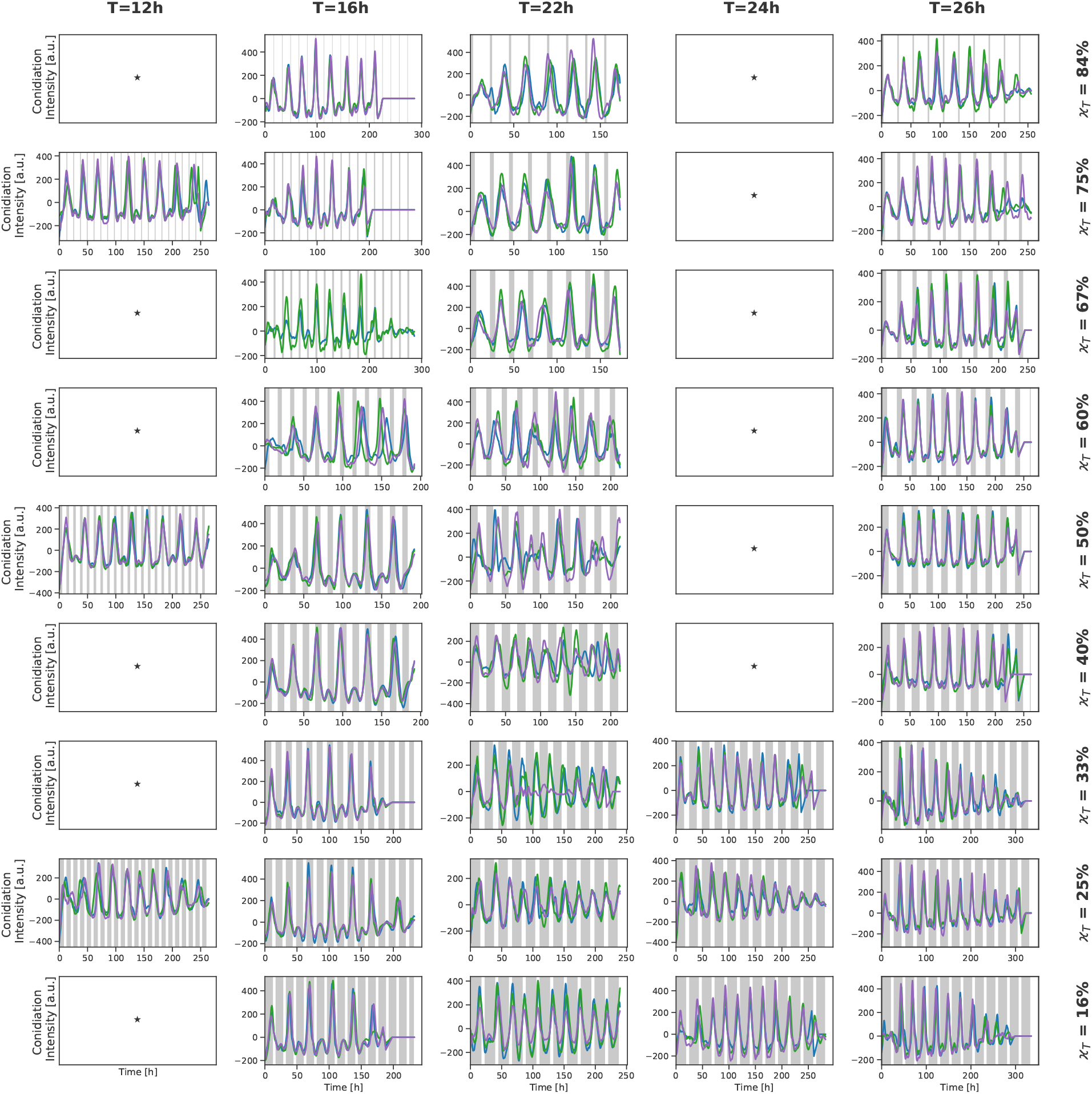
Experimental time series of the *frq* ^7^ strain. Same as Figure S1 for the *frq* ^7^ strain which has a free-running period of *τ_DD_* ≈ 29h under constant darkness and constant temperature.

**Figure S5:**
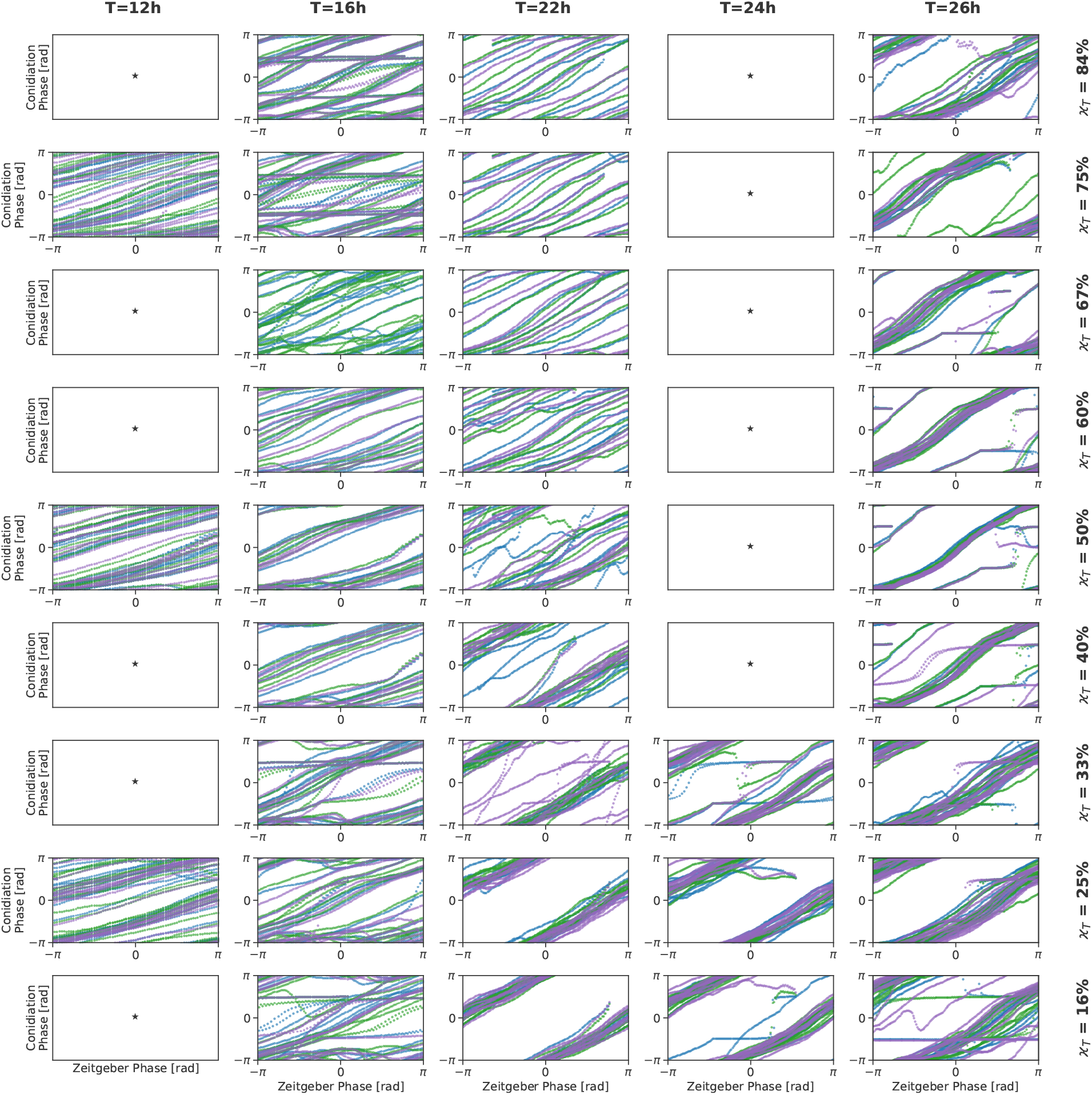
Phase plots for the *frq* ^7^ strain data. Same as Figure S2 for the *frq* ^7^ strain which has a free-running period of *τ_DD_* ≈ 29h under constant darkness and constant temperature.

**Figure S6:**
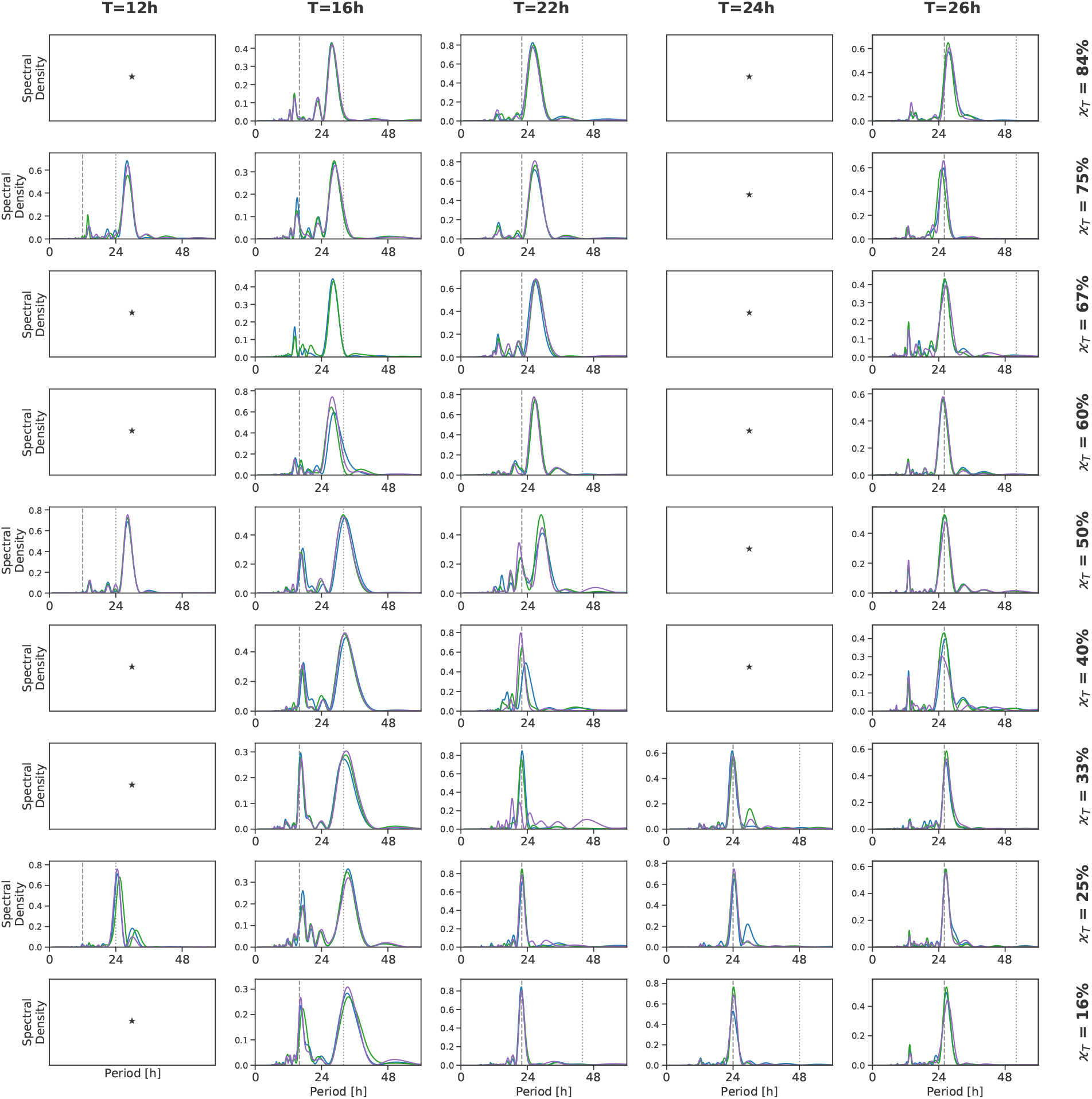
Lomb-Scargle periodograms for the *frq* ^7^ strain data. Same as Figure S2 for the *frq* ^7^ strain which has a free-running period of *τ_DD_* ≈ 29h under constant darkness and constant temperature.

**Figure S7:**
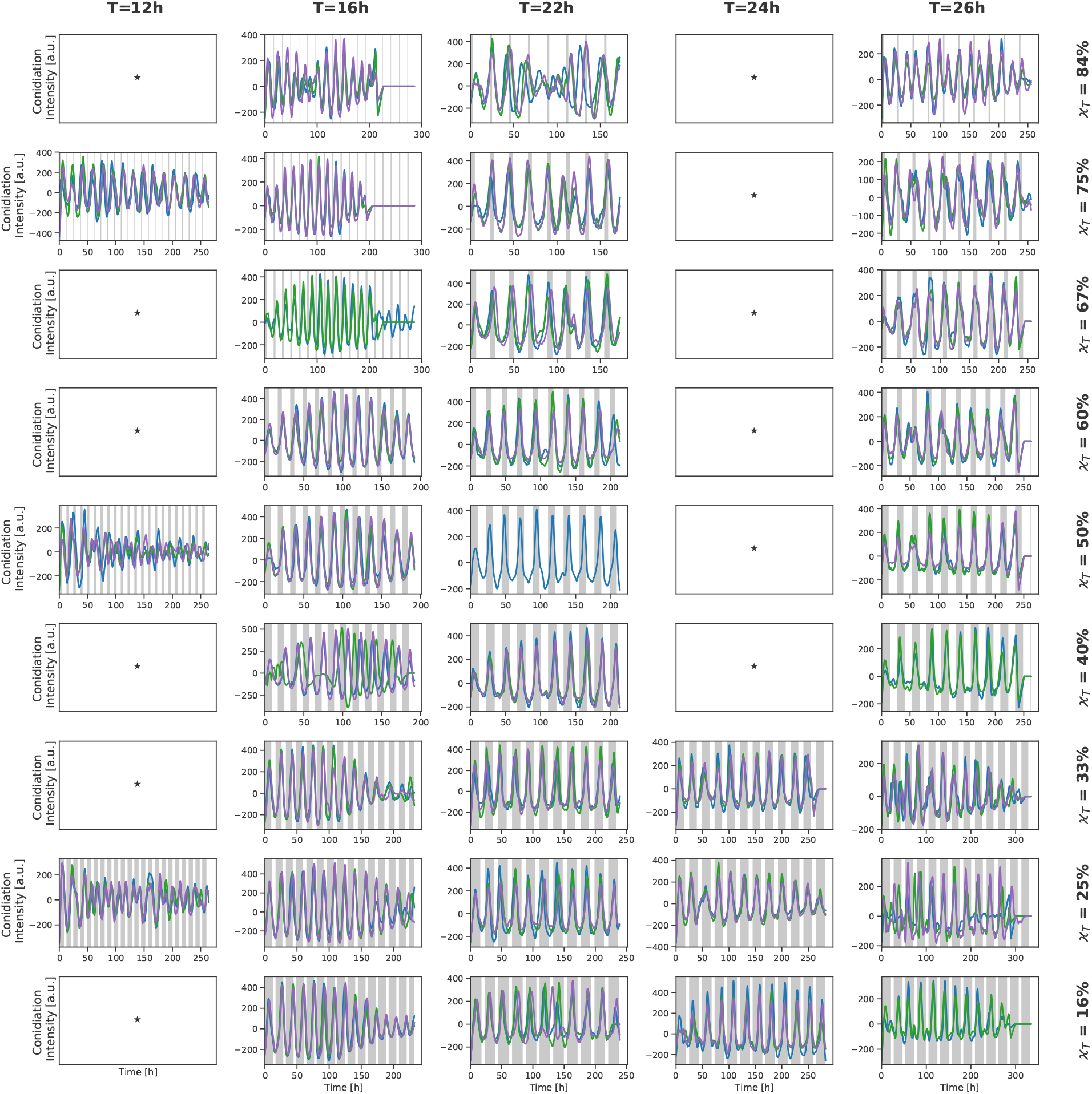
Experimental time series of the *frq* ^1^ strain. Same as Figure S1 for the *frq* ^1^ strain which has a free-running period of *τ_DD_* ≈ 16.5h under constant darkness and constant temperature.

**Figure S8:**
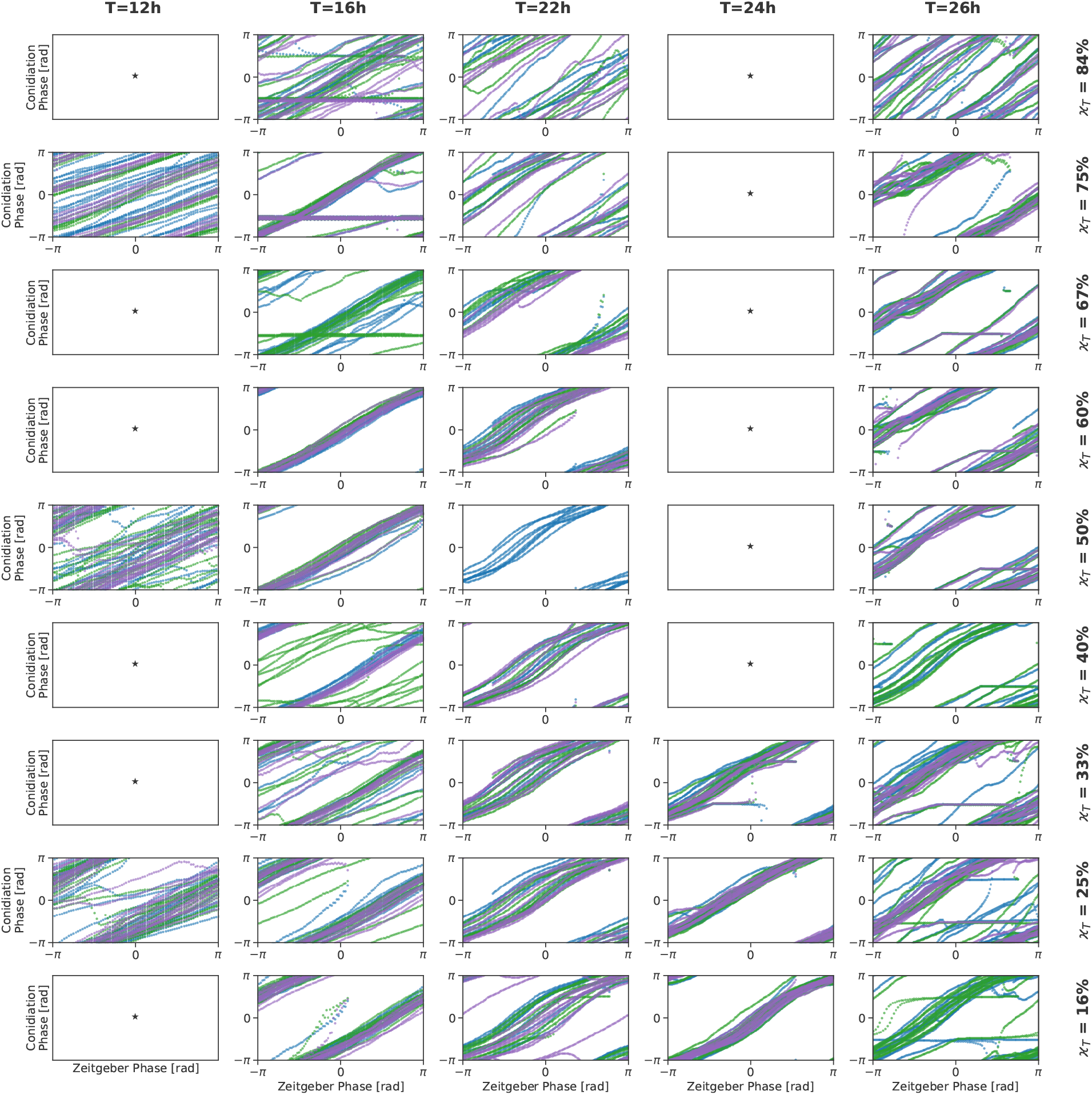
Phase plots for the *frq* ^1^ strain data. Same as Figure S2 for the *frq* ^1^ strain which has a free-running period of *τ_DD_* ≈ 16.5h under constant darkness and constant temperature.

**Figure S9:**
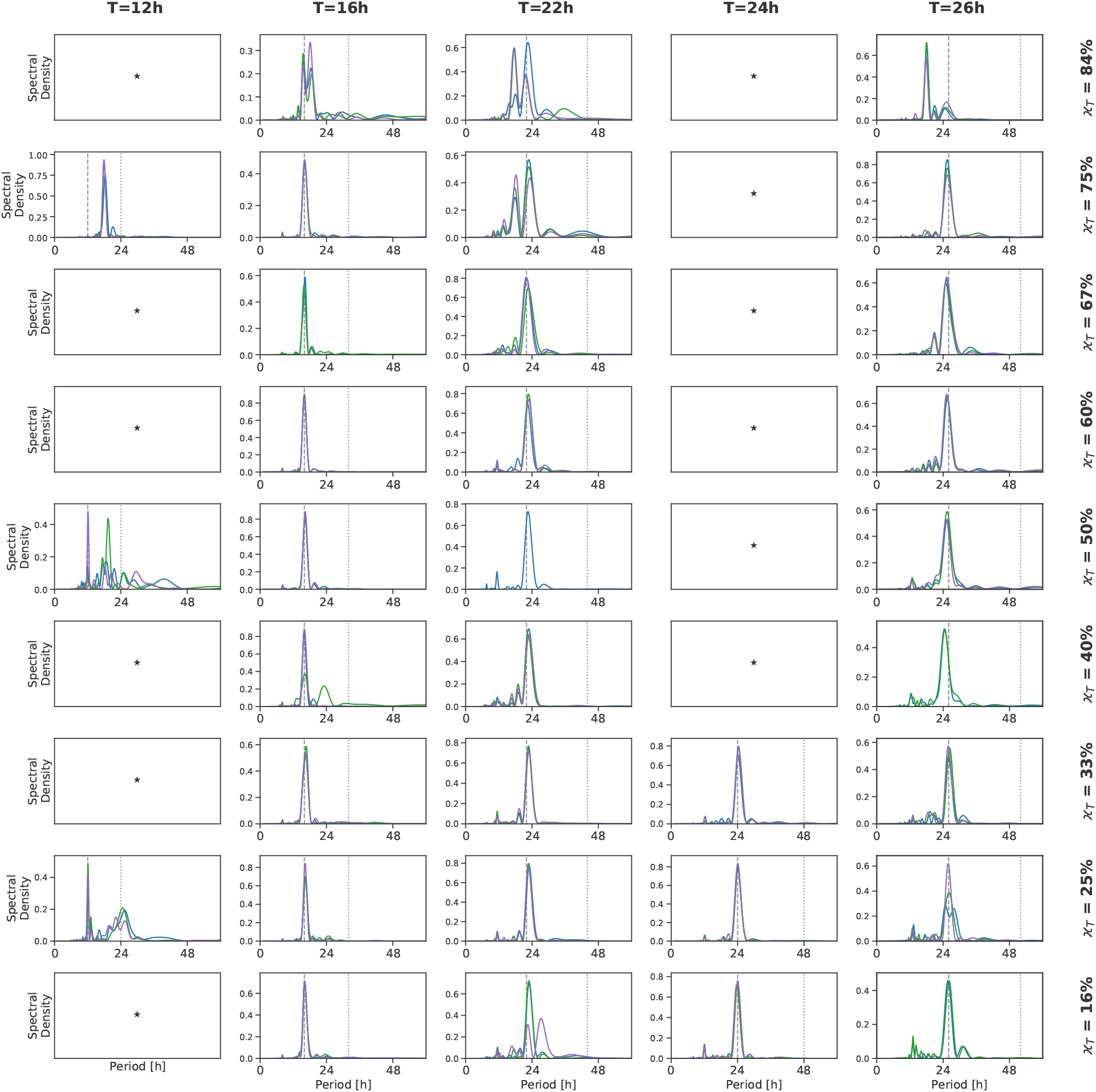
Lomb-Scargle periodograms for the *frq* ^1^ strain data. Same as Figure S3 for the *frq* ^1^ strain which has a free-running period of *τ_DD_* ≈ 16.5h under constant darkness and constant temperature.

**Figure S10:**
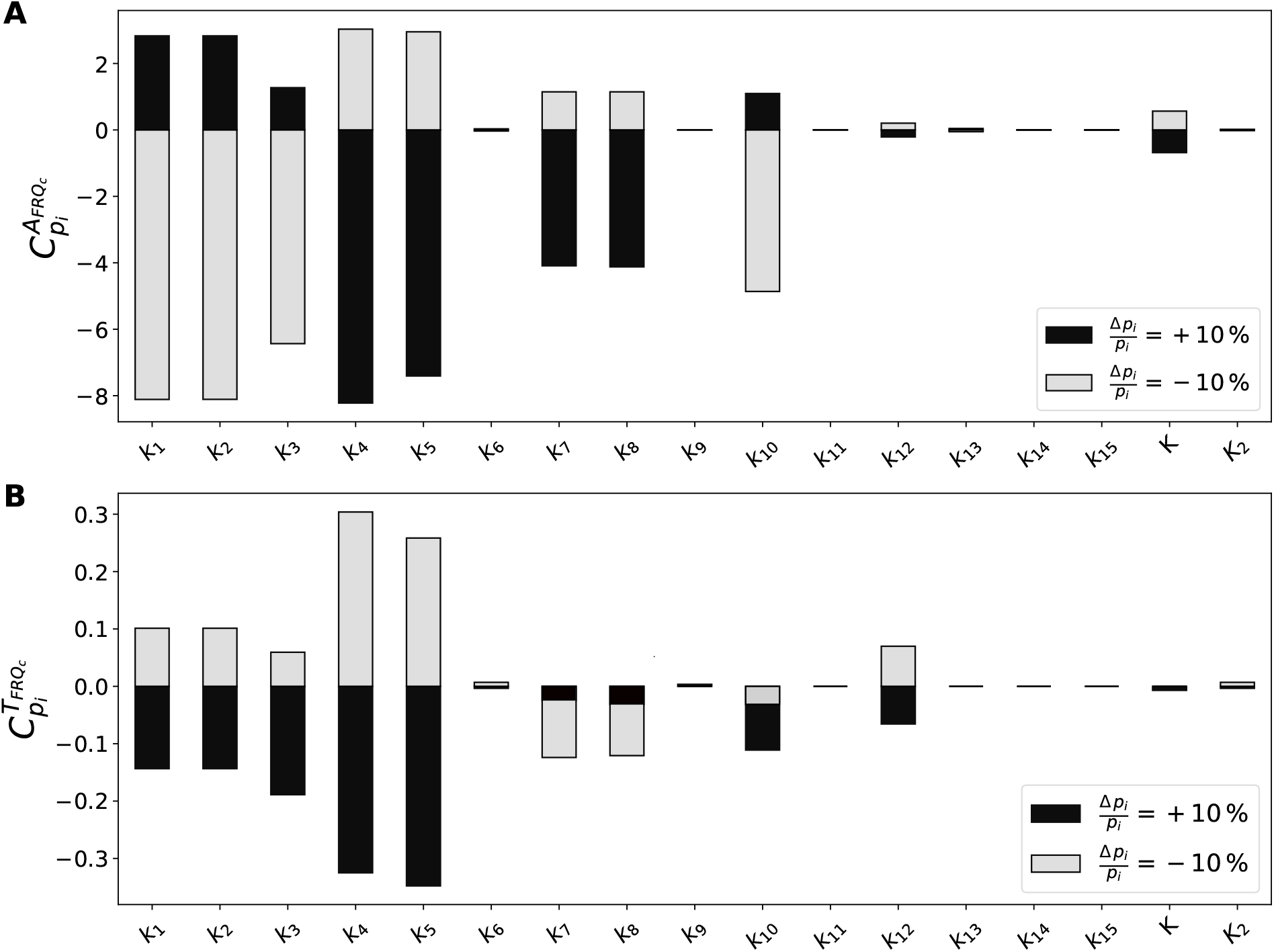
Sensitivity analysis of the Hong model. Control coefficients *C_p_i* for the relative change in FRQc amplitude (A) and FRQc period (B) for parameter variations of ±10 % (black and grey bars respectively). A sensitivity coefficient of 1 corresponds to a relative change of 10 %.

**Figure S11:**
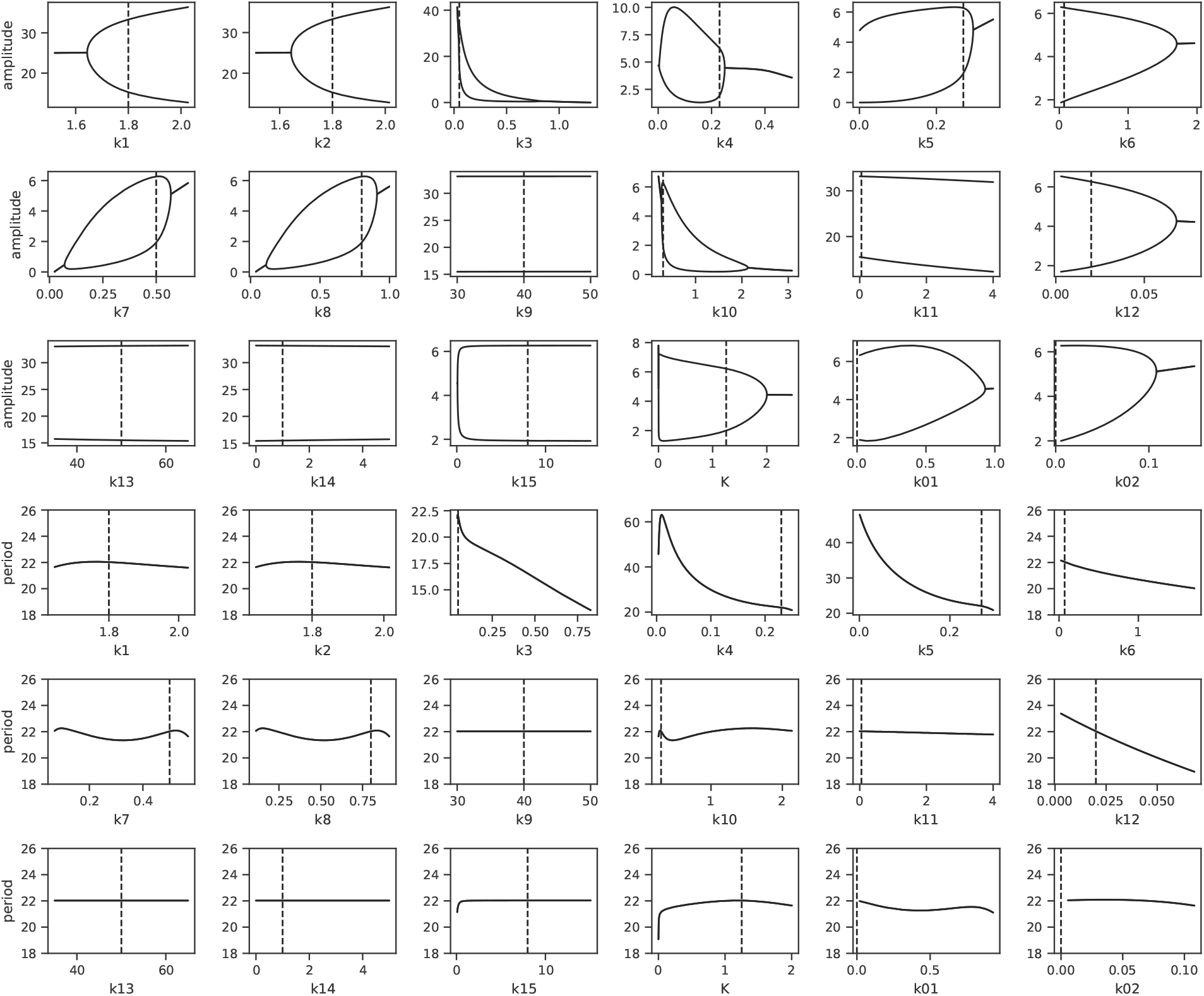
Bifurcation diagrams for all parameters of the Hong model. Parameters (x-axis) are plotted against either FRQc amplitude or FRQc period (y-axis) as indicated. Bifurcation analysis was performed using XPP-Auto.

**Figure S12:**
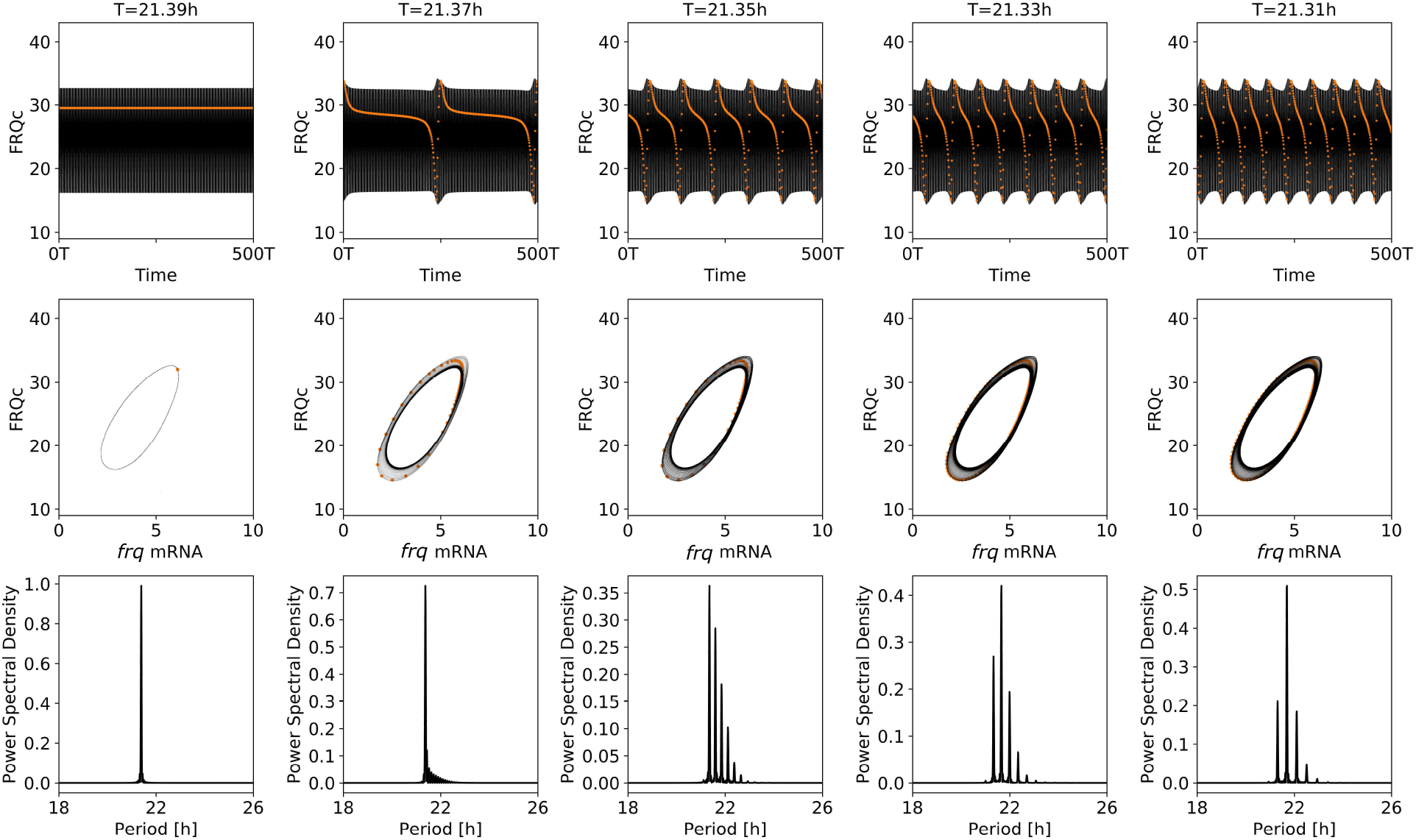
Leaving the entrainment regime at relatively small Zeitgeber strength typically results in modulations. Simulated dynamics (upper row), projection onto the two-dimensional phaseplane given by *frq* mRNA and FRQc concentrations (middle row) as well as Lomb-Scargle periodograms (bottom row). *Orange dots* indicate the corresponding stroboscopic map. Within the synchronization regime, the Lomb-Scargle spectrum contains a single peak at the period of the Zeitgeber signal as well as a stable fixed point for the stroboscopic map, i.e. a single point within the phase plane representation (left column). After leaving the synchronization regime, amplitude modulations occur (upper row), transforming the stroboscopic map into a closed curve (middle row) and leading to the formation of multiple peaks within the periodogram (bottom curve). Typical for amplitude modulations, the period of the amplitude envelope gets longer while leaving the diameter of the stroboscopic map approximately constant with an increasing distance from the synchronization regime. Simulations have been obtained for the nominal parameter set of the *Hong model* (see *main text*), *ϰ* = 0.75 and *z*_0_ = 0.04.

**Figure S13:**
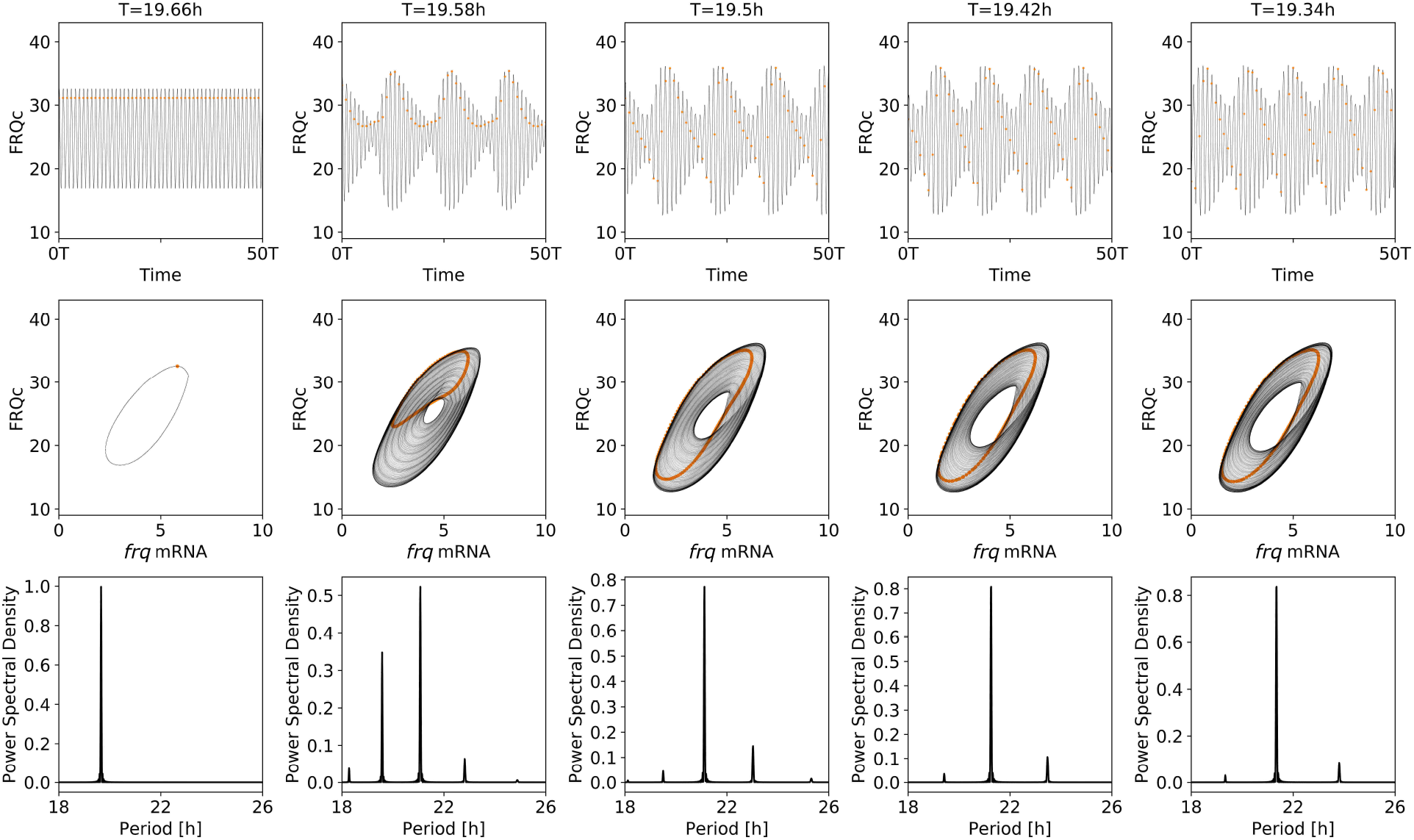
Leaving the entrainment regime at relatively high Zeitgeber strength typically results in beating. At relatively large Zeitgeber strength, i.e. within the upper parts of an *Arnold tongue*, the entrainment regime is typically left through a torus bifurcation. Similar to Fig. S12, simulated time series (upper row), a corresponding phase space representation (middle row) as well as Lomb-Sargle periodograms (bottom row) are shown. Simulations have been obtained for the nominal parameter set of the *Hong model* as in Fig. S12, *ϰ* = 0.75 and *z*_0_ = 0.2.

**Figure S14:**
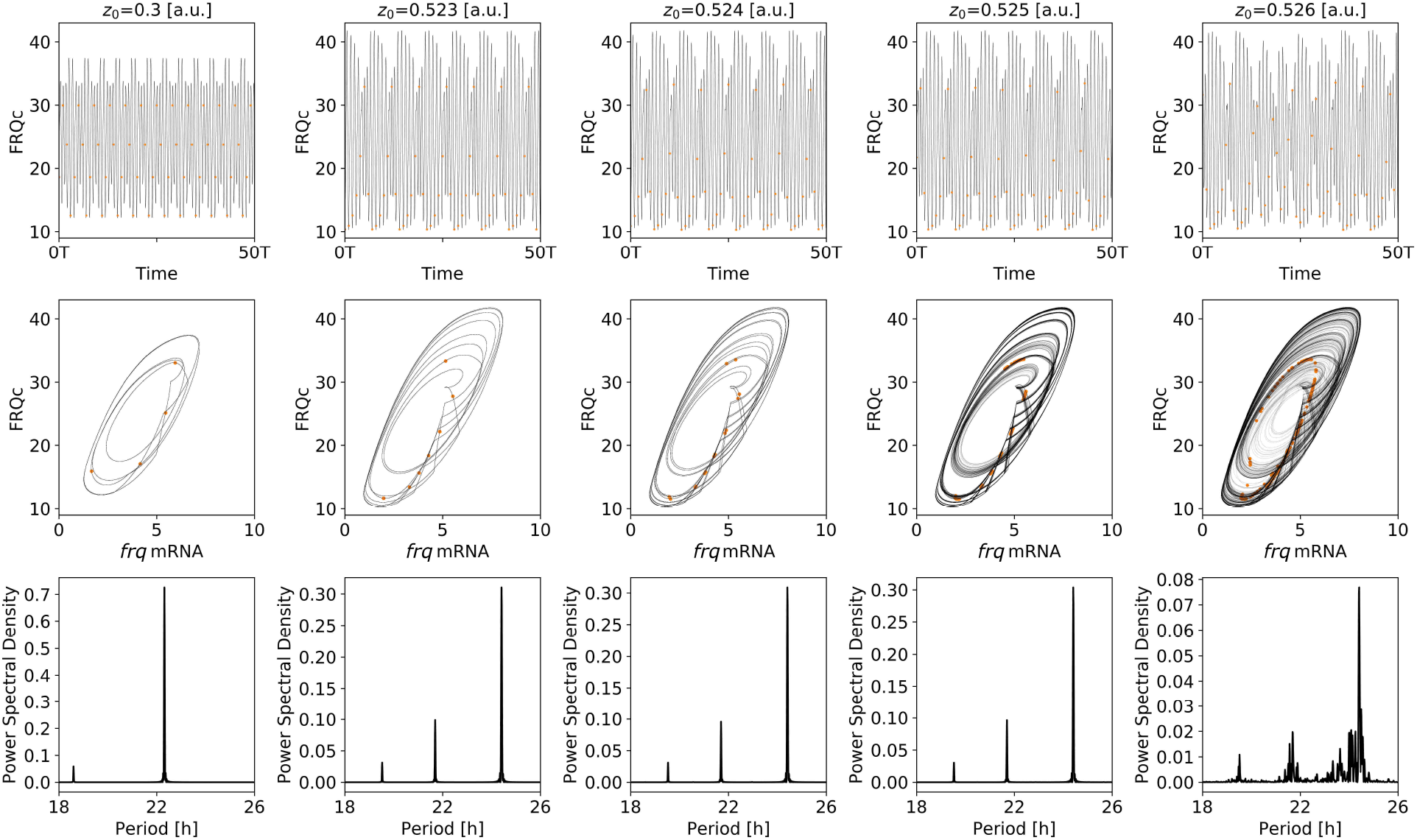
Route to chaos. The system undergoes various states of higher order synchronization (1st, 2nd and 3rd column) and shows chaotic dynamics for *z*_0_ = 0.526 (5th column) characterized by aperiodic, randomly fluctuating time series (5th column, upper row), a wrinkled aperiodic stroboscopic map (5th column, middle row, orange dots) and a continuous spectrum around the fundamental frequency and its harmonics (5th column, bottom row). Note the contraction of points in the stroboscopic map at the transition to chaos, indicating the higher order synchronized state for a slightly lower Zeitgeber strength (4th column, middle row, orange dots). Simulations for different Zeitgeber strength *z*_0_ have been obtained for the nominal parameter set of the *Hong model* (see *main text*), *ϰ* = 0.75 and *T* = 27.9h.

**Figure S15:**
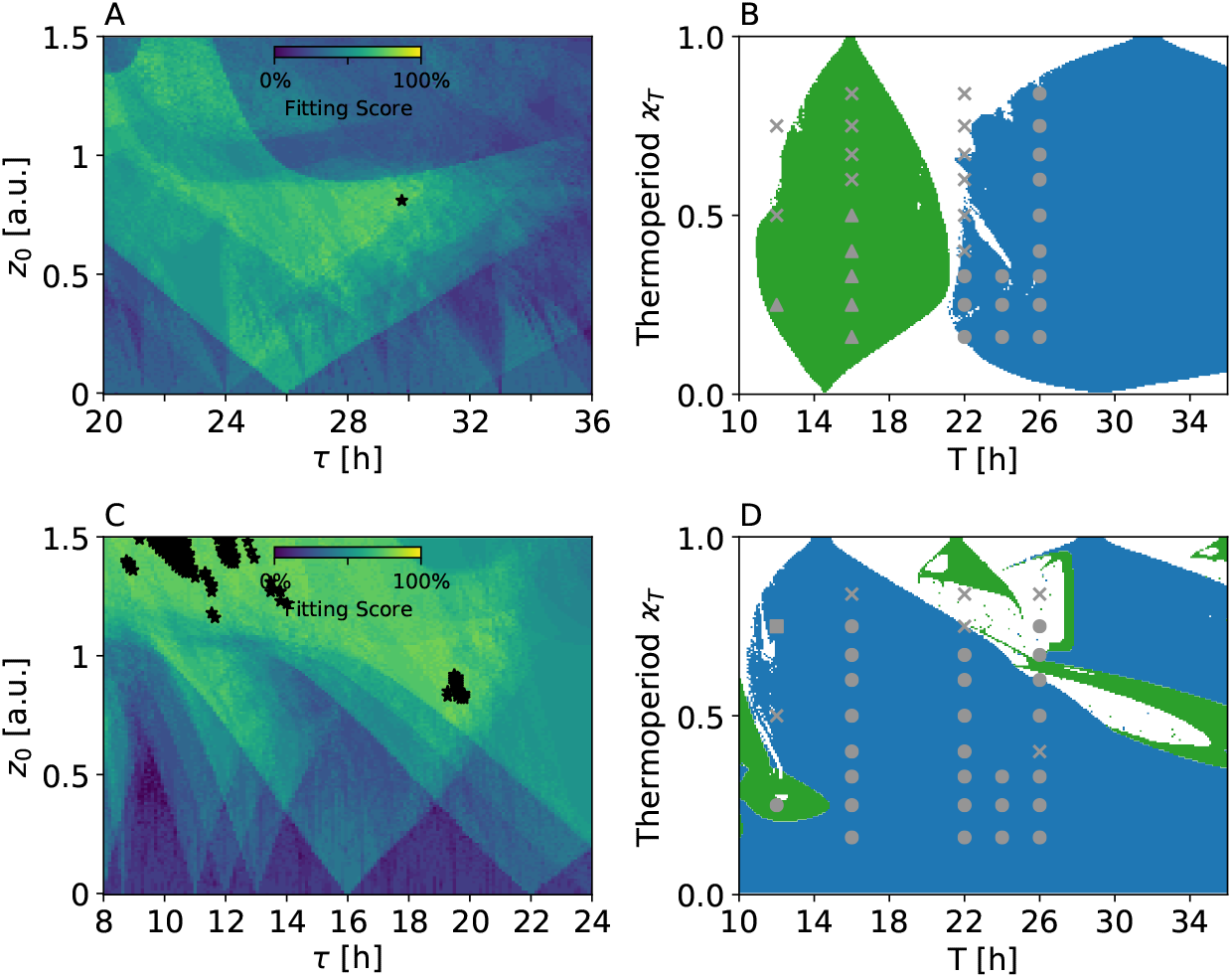
Experimentally observed entrainment behavior of *frq* ^7^ and *frq*^1^ mutant can be partially reproduced, using the previously published nominal parameter sets. A) Fitting scores for different Zeitgeber strength *z*_0_ and scaled internal free-running periods *τ* in case of the *frq*^7^ mutant, similar to Fig. 5 B of the *main text*. B) Experimentally observed dynamics (gray symbols) together with simulated 1:1 (*blue*) and 1:2 (*green*; frequency demultiplication) synchronization regions in the thermoperiod - Zeitgeber period parameter plane (Arnold onions). A Zeitgeber strength of *z*_0_ = 0.81 and scaled free-running period of *τ* ≈ 29.77h, corresponding to the *black star* in panel (A), has been used. C) Same as panel (A) for the *frq*^1^ mutant. D) Same as panel (B) for the *frq*^1^ mutant. A Zeitgeber strength of *z*_0_ = 1.22 and scaled free-running period of *τ* ≈ 14.01h has been used. *Stars* denote parameter combinations that lead to a maximal fitting score of ≈ 82% or 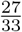 in case of both the *frq*^7^ (A) and *frq*^1^ (C) mutant. Parameter values used for simulations of mutant dynamics are described in *materials and methods* of the *main text*.

